# The universal suppressor mutation in the HSV-1 nuclear egress complex restores membrane budding defects by stabilizing the oligomeric lattice

**DOI:** 10.1101/2023.06.22.546118

**Authors:** Elizabeth B. Draganova, Hui Wang, Melanie Wu, Shiqing Liao, Amber Vu, Gonzalo L. Gonzalez-Del Pino, Z. Hong Zhou, Richard J. Roller, Ekaterina E. Heldwein

## Abstract

Nuclear egress is an essential process in herpesviral replication whereby nascent capsids translocate from the nucleus to the cytoplasm. This initial step of nuclear egress – budding at the inner nuclear membrane – is coordinated by the nuclear egress complex (NEC). Composed of the viral proteins UL31 and UL34, NEC deforms the membrane around the capsid as the latter buds into the perinuclear space. NEC oligomerization into a hexagonal membrane-bound lattice is essential for budding because mutations designed to perturb lattice interfaces reduce its budding ability. Previously, we identified an NEC suppressor mutation capable of restoring budding to a mutant with a weakened hexagonal lattice. Here, we show that the suppressor mutation can restore budding to a broad range of budding-deficient NEC mutants thereby acting as a universal suppressor. We demonstrate that the suppressor mutation indirectly promotes the formation of new contacts between the NEC hexamers that, ostensibly, stabilize the hexagonal lattice. This stabilization strategy is powerful enough to override the otherwise deleterious effects of mutations that destabilize the NEC lattice by different mechanisms, resulting in a functional NEC hexagonal lattice and restoration of membrane budding.

## INTRODUCTION

Viruses are experts at reorganizing host membranes to traffic their capsids across the compartmentalized interior of eukaryotic cells. One of the more unusual mechanisms of membrane manipulation is found in *Herpesvirales*, which is an order of large, enveloped viruses that infect multiple species across the animal kingdom and cause life-long infections in the majority of the world’s population. Replication of herpesviral dsDNA genomes and their subsequent packaging into capsids occurs within the nucleus. Genome-containing capsids are then transported into the cytoplasm for maturation into infectious virions. Most of the nucleocytoplasmic traffic occurs through the nuclear pores, but at ∼125 nm in diameter, herpesviral capsids are too large to fit through the ∼40-nm nuclear pore opening. Instead, the capsids use a complex, non-canonical nuclear transport route termed nuclear egress ^1–3^. First, they dock at the inner nuclear membrane (INM) and bud into the perinuclear space, producing perinuclear enveloped virions (a stage termed primary envelopment). The envelopes of these intermediates then fuse with the outer nuclear membrane (ONM) and capsids are then released into the cytoplasm (a stage termed de-envelopment).

The nuclear egress mechanism is best understood for the family *Herpesviridae*, commonly referred to as herpesviruses, which infect mammals, birds, and reptiles. Two conserved viral proteins, called UL31 and UL34 in herpes simplex virus (HSV), are essential for nuclear egress in herpesviruses. UL31 is a soluble nuclear phosphoprotein ^4,5^ whereas UL34 is a type I membrane protein containing a single C-terminal transmembrane helix ^4,6^. Together, UL31 and UL34 form the heterodimeric nuclear egress complex (NEC) that is anchored in the INM and faces the nuclear interior. Both proteins are essential for nuclear egress, and in the absence of either, capsids become trapped within the nucleus, which results in greatly reduced viral titers ^4,5, 7–11^. Moreover, overexpression of both UL31 and UL34 in uninfected cells causes the accumulation of empty budded vesicles in the perinuclear space, which implies that the NEC is not only necessary but also sufficient for the INM budding ^12–16^. Collectively, these findings highlight the central role of the NEC during nuclear egress.

Recent studies with purified recombinant NEC and synthetic lipid vesicles have shown that several NEC homologs can deform and bud membranes *in vitro* in the absence of added energy or other proteins. These include the NECs from herpes simplex virus 1 (HSV-1) ^17^, a prototypical herpesvirus that infects much of the world’s population; the closely related pseudorabies virus (PRV) that infects animals ^18^; and the more distantly related Epstein-Barr Virus (EBV) ^19^, a nearly ubiquitous human herpesvirus.

The NEC oligomerizes into membrane-bound coats on the inner surface of the budded vesicles. Hexagonal coats resembling a honeycomb have been observed by cryo-electron microscopy/tomography (cryo-EM/ET) on vesicles formed by recombinant HSV-1 NEC *in vitro* ^17^, vesicles formed in uninfected cells overexpressing PRV NEC ^20^, and in perinuclear vesicles formed in HSV-1-infected cells ^21^. Interestingly, crystallized NEC homologs from HSV-1 ^22^ and human cytomegalovirus (HCMV) ^23^ also formed hexagonal crystal lattices of geometry and dimensions similar to those observed in the membrane-bound NEC coats. Finally, EBV NEC also forms membrane-bound coats *in vitro* but their geometry is yet unclear ^19^. Both the intrinsic membrane budding ability and the formation of oligomeric coats thus appear to be conserved among the NEC homologs.

In HSV-1 NEC, oligomerization into the hexagonal lattice is essential for budding. Mutations targeting lattice interfaces within the NEC hexamers (hexameric) or between hexamers (interhexameric) cause budding defects *in vitro* ^17,22^ and reduce nuclear egress in infected cells ^24,25^. The first such mutation, D35A_UL34_/E37A_UL34_, was identified in a mutational screen targeting charge clusters in the HSV-1 UL34 sequence ^26^. This double mutation reduced viral titers by ∼3 orders of magnitude to levels of UL34-null mutant HSV-1 and blocked capsid egress from the nucleus in a dominant-negative manner ^25^. Therefore, we refer to it as DN_UL34_. The mutation did not affect the NEC formation, its localization to the INM, or capsid docking at the INM but, instead, precluded capsid budding ^25^. Furthermore, purified recombinant NEC-DN_UL34_ bound synthetic membranes *in vitro* but had minimal membrane-budding activity and did not form hexagonal coats on membranes ^17^.

Interestingly, the nuclear budding defect due to the DN_UL34_ mutation could be suppressed by an extragenic mutation in HSV-1 UL31, R229L_UL31_, which arose during serial passaging of the DN_UL34_ mutant HSV-1 virus on a UL34-complementing cell line ^25^. We refer to this mutation as SUP_UL31_. SUP_UL31_ maps near the interhexameric interface, far away from the DN_UL34_ mutations at hexameric interface ^22^, making it unclear how the SUP_UL31_ mutation restores DN_UL34_ nuclear budding defects.

Here, we show that the SUP_UL31_ mutation can restore efficient budding to a broad range of mutants that disrupt important functional interfaces, acting as a “universal” suppressor of budding defects. Using cryo-ET and x-ray crystallography, we show that the SUP_UL31_ mutation does not change the structure of the NEC heterodimer or its oligomerization into hexamers. Instead, it promotes the formation of new contacts at the interhexameric interface. We propose that the increased interhexameric interface reinforces the hexagonal NEC lattice, thereby counteracting the deleterious effects of mutations that perturb it.

## RESULTS

### The SUP_UL31_ mutation restores membrane budding in vitro to various oligomeric interface mutants

HSV-1 NEC oligomerizes into a hexagonal lattice (**Fig. 1a**) stabilized by interactions between NEC heterodimers within hexamers (hexameric interface; **Fig. 1b**) and between hexamers (interhexameric interface; (**Fig. 1c**). NEC hexameric lattice formation is essential for membrane budding because mutations engineered to disrupt lattice interfaces reduce budding *in vitro* ^17,22^ and nuclear egress in infected cells ^24,25^. These budding-deficient mutations are D35A_UL34_/E37A_UL34_ (DN_UL34_), V92F_UL34_, T123Q_UL34_, V247F_UL31_, and F252Y_UL31_ at the hexameric interface (**Fig. 1b**) and E153R_UL31_ at the interhexameric interface **(Fig. 1c)**. The SUP_UL31_ mutation restores budding *in vitro* to DN_UL34_ and V92F_UL34_ mutants ^22^. Here, we asked if it could restore budding to other interface mutants, T123Q_UL34_ and F252Y_UL31_ (hexameric) **(Fig. 1b)** and E153R_UL31_ (interhexameric) **(Fig. 1c)**.

**Fig. 1.**
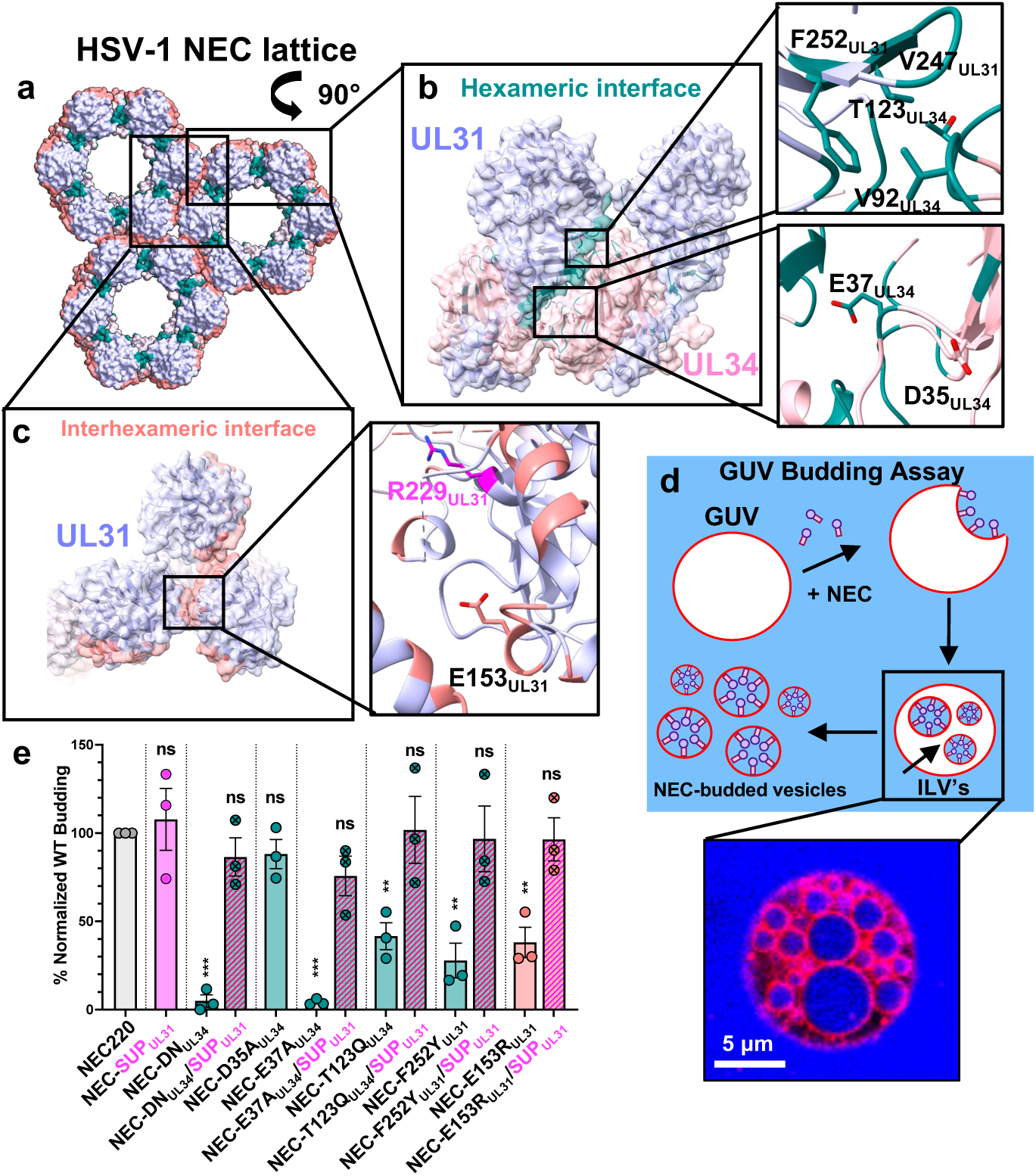
The SUP_UL31_ mutation restores budding activity to budding-deficient oligomeric interface NEC mutants. **a-c)** HSV-1 NEC hexameric and interhexameric interfaces highlighting the locations of residues mutated for this study. **d)** A cartoon representation of the GUV budding assay showing the NEC (purple circles and pink rectangles) binding to red fluorescent GUVs and undergoing negative curvature to form an NEC-coated intraluminal vesicle (ILV). Free NEC continues to bud the GUVs until only fully-budded vesicles containing NEC on the interior remain. Cascade blue, a membrane impermeant dye, is used to monitor budding. **e)** SUP-R229L_31_ rescues budding in both hexameric and interhexameric budding-deficient NEC mutants *in vitro*. The percentage of *in vitro* budding was determined by counting the number of ILVs within the GUVs after the addition of NEC220 or the corresponding NEC mutant. A background count, the number of ILVs in the absence of NEC, was subtracted from each condition. Each construct was tested in three biological replicates, each consisting of three technical replicates. Symbols show the average budding efficiency of three technical replicates compared to NEC220 (100%; grey). Significance was calculated using an unpaired Student’s t-test with Welch’s correction (P < 0.001 = ***; P < 0.0001 = **; ns = not significant) in GraphPad Prism 9.0.

We also wanted to assess the individual contributions of D35A_UL34_ and E37A_UL34_ mutations to the budding-deficient phenotype of the DN_UL34_ mutant. Residue E37_UL34_ is located at the hexameric interface where its side chain forms a hydrogen bond with T89_UL31_ of the neighboring NEC heterodimer. The E37A_UL34_ mutation eliminates this hydrogen bond, which would disrupt the hexameric interface. Indeed, the E37A_UL34_ mutant was deficient in budding *in vitro* ^22^. However, the side chain of residue D35_UL34_ points away from the hexameric interface (**Fig. 1b**). Therefore, we tested if the D35A_UL34_ mutation would have any effect on budding.

The *in-vitro* budding activity of all NEC mutants was measured by an established assay ^17,22,27,28^ utilizing fluorescently labeled giant unilamellar vesicles (GUV), soluble fluorescent dye Cascade Blue, and the soluble version of HSV-1 NEC, NEC220, which contained full-length UL31 and UL34 residues 1-220 **(Fig. 1d)**. We first confirmed the *in-vitro* phenotypes of the budding-deficient mutants. Both DN_UL34_ and E37A_UL34_ mutations reduced budding to ∼10% of the WT NEC220 **(Fig. 1e),** consistent with our previous findings ^22^ whereas the D35A_UL34_ mutation alone had no effect **(Fig. 1e).** Thus, the E37A_UL34_ mutation is solely responsible for the nonbudding phenotype of DN_UL34_. The interface mutations T123Q_UL34_, F252Y_UL31_, and E153R_UL31_ (**Fig. 1bc**) reduced budding to ∼30-40% of the WT NEC220 **(Fig. 1e)**, as previously observed ^22^. The SUP_UL31_ mutation did not affect the budding efficiency of the WT NEC220 but restored budding not only to the DN_UL34_ as shown previously ^22^ but to all other lattice interface mutants regardless of their location **(Fig. 1e)**. Thus, the SUP_UL31_ mutation can restore efficient budding to a broad range of lattice interface mutants.

### The SUP_UL31_ mutation complements the growth defects of HSV-1 containing oligomeric interface mutations

To correlate the *in-vitro* budding phenotypes with the infected cell phenotypes, we used an established viral growth complementation assay ^9^. This assay measures the ability of a mutant protein expressed *in trans* to complement the poor growth of a virus lacking the corresponding gene (the so-called null virus). Hep-2 cells were transfected with plasmids encoding either WT, mutant UL34 (D35A_UL34_/E37A_UL34_, D35A_UL34_, E37A_UL34_, T123Q_UL34_), or mutant UL31 (F252Y_UL31_ and E153R_UL31_) and then infected with either a UL34-null HSV-1 or UL31-null HSV-1. The amount of infectious viral progeny produced was measured by plaque assay on either UL34-expressing **(Fig. 2a)** or UL31-expressing Vero cells **(Fig. 2b).** We found that cells expressing the D35A_UL34_/E37A_UL34_, E37A_UL34_, T123Q_UL34_, or E153R_UL31_ mutants poorly complemented replication of either the UL34-null **(Fig. 2a)** or UL31-null HSV-1 **(Fig. 2b)**, respectively, *in trans*, in agreement with their impaired budding phenotypes *in vitro*. By contrast, the D35A_UL34_ mutant complemented UL34-null HSV-1 *in trans* with an efficiency similar to that of the WT UL34 **(Fig. 2a)**. Surprisingly, the F252Y_UL31_ mutant complemented UL31-null HSV-1 *in trans* similarly to WT UL31 despite reduced budding efficiency in our *in-vitro* budding assay **(Fig. 1e)**.

**Fig. 2.**
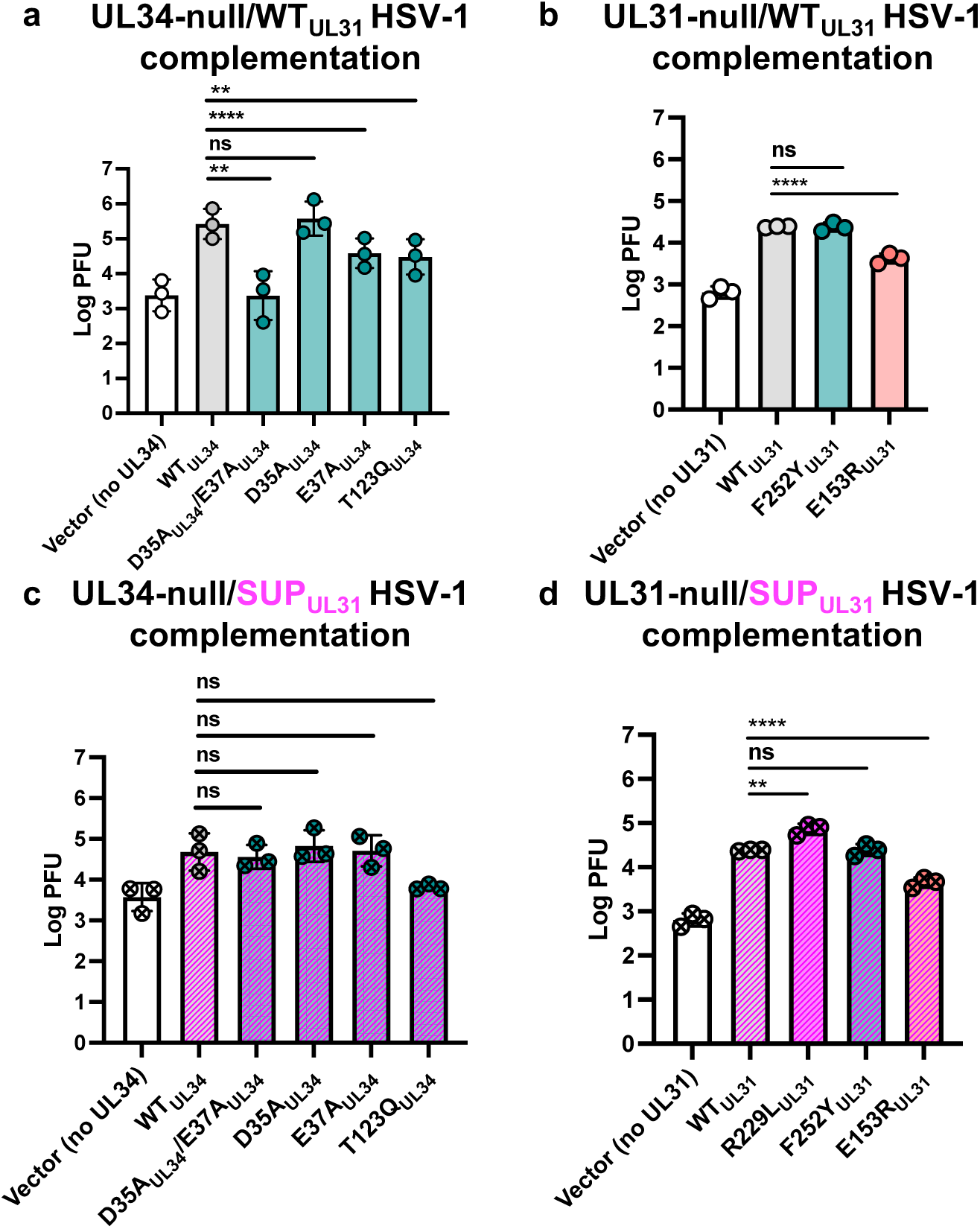
The SUP_UL31_ mutation complements the growth defects of HSV-1 containing oligomeric interface mutations. **(a-b)** WT_UL31_ can only complement the growth of the D35A_UL34_ and F252Y_UL31_ oligomeric interface mutants whereas SUP_UL31_ can complement more **(c-d)**. For all experiments, Hep-2 cells were transfected with the corresponding UL34 or UL31 mutant plasmid and infected with either a UL34-null/UL31_WT_ virus **(a),** a UL31-null/UL31_WT_ virus **(b)**, a UL34-null/UL31_R229L_ virus **(c)**, or a UL31-null/UL31_R229L_ virus **(d)**. Each bar represents the mean of three independent experiments. Statistical significance was determined by performing a paired two-tailed t-test of each mutant against the WT in GraphPad Prism. P < 0.01 = **; P < 0.0001 = ****; ns, not significant.

To rule out increased protein expression levels as the *in-trans* complementation mechanism, we measured expression levels of transfected WT UL34, WT UL31, and the corresponding mutant proteins during infection with the corresponding null virus. All mutant UL34 proteins expressed at levels similar to that of WT UL34 **(Fig. S1)**. Reduced complementation efficiencies of D35A_UL34_/E37A_UL34_, E37A_UL34_, and T123Q_UL34_ (**Fig. 2a**) are thus due to the specific mutation(s). In contrast, both F252Y_UL31_ and E153R_UL31_ are overexpressed relative to the WT UL31 and R229L_UL31_. Efficient complementation by the F252Y_UL31_ mutant is likely due to its higher expression levels (**Fig. 2b**).

To probe the ability of the SUP_UL31_ mutation to restore efficient complementation to the UL34 and UL31 mutants, i.e., to suppress their complementation defects, we generated the UL34-null/SUP_UL31_ and UL31-null/SUP_UL31_ mutant HSV-1 viruses. Hep-2 cells were transfected with plasmids encoding either WT UL34, mutant UL34 (D35A_UL34_/E37A_UL34_, D35A_UL34_, E37A_UL34_, T123Q_UL34_), WT UL31, or mutant UL31 (F252Y_UL31_ and E153R_UL31_), and then infected with either a UL34-null/SUP_UL31_ or UL31-null/SUP_UL31_ HSV-1 (instead of a UL34-null or UL31-null HSV-1). HSV-1 containing the SUP_UL31_ mutation replicates less efficiently than the WT HSV-1, yielding ∼1-log-fold lower viral titer **(Fig. 2cd),** as reported previously ^25^. Therefore, complementation of either the UL34-null/SUP_UL31_ or UL31-null/SUP_UL31_ viruses by the WT UL34 or UL31, respectively, was used as a reference point for assessing the ability of the SUP_UL31_ mutation to restore efficient complementation to UL34 and UL31 mutants. Indeed, the SUP_UL31_ mutation suppressed the complementation defects of both the D35A_UL34_/E37A_UL34_ and E37A_UL34_ mutants similarly to the WT UL34 **(Fig. 2c).** Unsurprisingly, the SUP_UL31_ mutation had no obvious effect on the already efficient complementation by the D35A_UL34_ **(Fig. 2c)** and F252Y_UL31_ mutants **(Fig. 2d).** However, it was unable to fully restore the poor complementation by either the T123Q_UL34_ **(Fig. 2c)** or E153R_UL31_ mutants **(Fig. 2d)** despite restoring their budding defects *in vitro.* We hypothesize that the T123Q_UL34_ and E153R_UL31_ mutations may impair some other important viral replication function of UL34 or UL31, respectively, that cannot be suppressed by the SUP_UL31_ mutation, e.g., nuclear lamina dissolution, capsid docking at the INM, or capsid recruitment.

### The SUP_UL31_ mutation restores efficient budding in vitro to heterodimeric interface mutants and complements their viral growth defects

The aforementioned mutational screen targeting charge clusters in the HSV-1 UL34 sequence ^26^, identified another double mutant, K137A_UL34_/R139A_UL34_, that could not trans-complement the growth of the HSV-1 UL34-null virus. This suggested that residues K137_UL34_ and R139_UL34_ are important for HSV-1 replication. The double mutation did not affect the NEC localization to the INM, suggesting a defect in the NEC function ^26^. In the HSV-1 NEC crystal structure, K137_UL34_ forms salt bridges with E67_UL34_ and Y61_UL31_ at the heterodimeric interface between the globular domains of UL31 and UL34 **(Fig. 3a, inset)**. Thus, K137_UL34_ could contribute to the stabilization of the NEC heterodimer. By contrast, R139_UL34_ does not form any obvious interactions **(Fig. 3a, inset)**.

**Fig. 3.**
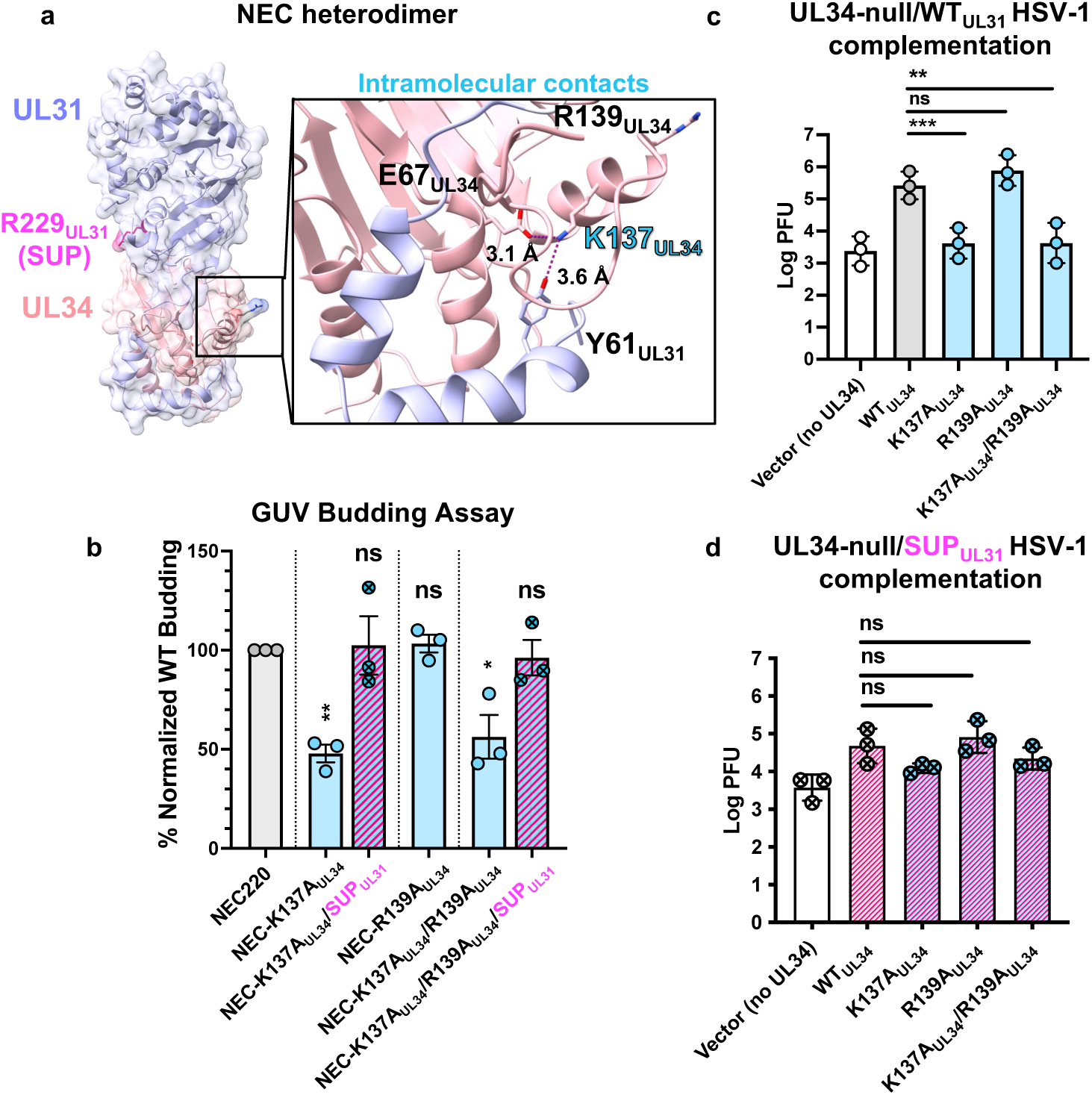
The SUP_UL31_ mutation restores budding to heterodimeric interface mutants and complements viral growth defects. **a)** Locations of intramolecular NEC residues mutated for this study. Inset shows interactions between various residues at the heterodimeric interface thought to be important for NEC heterodimer stabilization. **b)** NEC-SUP_UL31_ rescues budding of NEC heterodimeric interface mutants *in vitro*. The percentage of *in-vitro* budding was determined by counting the number of ILVs within the GUVs after the addition of NEC220 or the corresponding NEC mutant. A background count, the number of ILVs in the absence of NEC, was subtracted from each condition. Each construct was tested in three biological replicates, each consisting of three technical replicates. Symbols show the average budding efficiency of three technical replicates compared to NEC220 (100%; grey). Significance was calculated using an unpaired Student’s t-test with Welch’s correction (P < 0.05 = *; P < 0.01 = **; ns = not significant) in GraphPad Prism 9.0. **c)** WT_UL34_ can only complement the growth of the R139A_UL34_ heterodimeric interface mutant whereas SUP_UL31_ **(d)** can partially complement more. For both experiments, Hep-2 cells were transfected with the corresponding UL34 mutant plasmid and infected with a UL34-null/UL31 WT virus **(c)** or a UL34-null/UL31_R229L_ virus **(d)**. Each bar represents the mean of three independent experiments. Statistical significance was determined by performing a one-way ANOVA on log-converted values using the Method of Tukey for multiple comparisons implemented on GraphPad Prism. **, P < 0.01 = **; P < 0.001 = ***; ns, not significant.

To test the effect of the K137A_UL34_, R139A_UL34_, and K137A_UL34_/R139A_UL34_ mutations on the heterodimer stability and budding activity *in vitro*, we introduced them into the recombinant NEC220. Typically, size-exclusion chromatography on samples of purified, WT NEC220 yields only fractions containing equimolar amounts of UL31 and UL34, indicating the intact UL31:UL34=1:1 complex ^17^. Indeed, this pattern was observed for the NEC220-R139A_UL34_ mutant **(Fig. S2a).** However, both NEC220-K137A_UL34_ and NEC220-K137A_UL34_/R139A_UL34_ mutants also yielded fractions containing free UL34 or fractions containing more UL34 than UL31 **(Fig. S2bc)** despite equimolar amounts of UL31 and UL34 being loaded onto the size-exclusion column. Thus, the K137A_UL34_ mutation appeared to destabilize the NEC heterodimer. No free UL31 was detected in any of the fractions, suggesting that it may have been retained on a filter within the chromatography line. Only fractions containing equimolar amounts of UL31 and UL34 were used for further characterization.

Both K137A_UL34_ and K137A_UL34_/R139A_UL34_ mutations reduced budding to ∼50% of the WT NEC220 whereas the R139A_UL34_ mutation had no effect **(Fig. 3b)**. The K137A_UL34_ mutation is thus solely responsible for the defective budding phenotype of the double K137A_UL34_/R139A_UL34_ mutant. Surprisingly, the SUP_UL31_ mutation fully restored efficient budding to both K137A_UL34_ and K137A_UL34_/R139A_UL34_ mutants **(Fig. 3b).** But the mutant NEC heterodimers remained unstable **(Fig. S3)**. Therefore, the SUP_UL31_ mutation does not restore budding by restoring heterodimer stability.

To assess the effects of these mutations on viral replication, we performed the viral growth complementation assay described above. Both K137A_UL34_ and R139A _UL34_ proteins are expressed at levels similar to WT (**Fig. S1**). R139A_UL34_ complemented the growth of both the UL34-null **(Fig. 3c)** and UL34-null/SUP_UL31_ viruses on par with the WT UL34 **(Fig. 3d).** As expected, K137A_UL34_ and K137A_UL34_/R139A_UL34_ complemented growth of the UL34-null virus poorly **(Fig. 3c),** which is consistent with their *in-vitro* budding defects. However, both mutants complemented the growth of the UL34-null/SUP_UL31_ virus almost as efficiently as the WT UL34 **(Fig. 3d)**. Therefore, SUP_UL31_ mutation can restore both budding and replication defects caused by the K137A_UL34_ mutation.

### The SUP_UL31_ mutation partially restores budding in vitro to a membrane interface mutant

In addition to UL31/UL34 and NEC/NEC interfaces, the NEC/membrane interface is also functionally important in HSV-1 NEC. Both UL31 and UL34 contain membrane-proximal regions (MPRs) **(Fig. 4ab)** that mediate membrane association ^17,27^ and are essential for budding *in vitro* ^27^. The UL31 MPR contains clusters of positively charged residues that interact with model membranes and increase lipid order, which leads to membrane deformation and budding ^27^. The UL31 MPR also contains six serines **(Fig. 4b)** that are phosphorylated during infection ^5^ by the HSV-1 kinase US3 ^29^. Phosphomimicking serine-to-glutamate mutations of these six serines (SE6_UL31_) **(Fig. 4b)** reduce nuclear egress and viral titers during HSV-1 infection ^30^ and impair NEC/membrane interactions and budding activity *in vitro* ^27^. Previously, we proposed that negative charges introduced by phosphorylation or phosphomimicking mutations reduce electrostatic interactions between the MPR and the lipid headgroups that are necessary for membrane deformation and budding ^27^. Here, we asked whether the SUP_UL31_ mutation could restore budding *in vitro* to the budding-deficient SE6_UL31_ mutant.

**Fig. 4.**
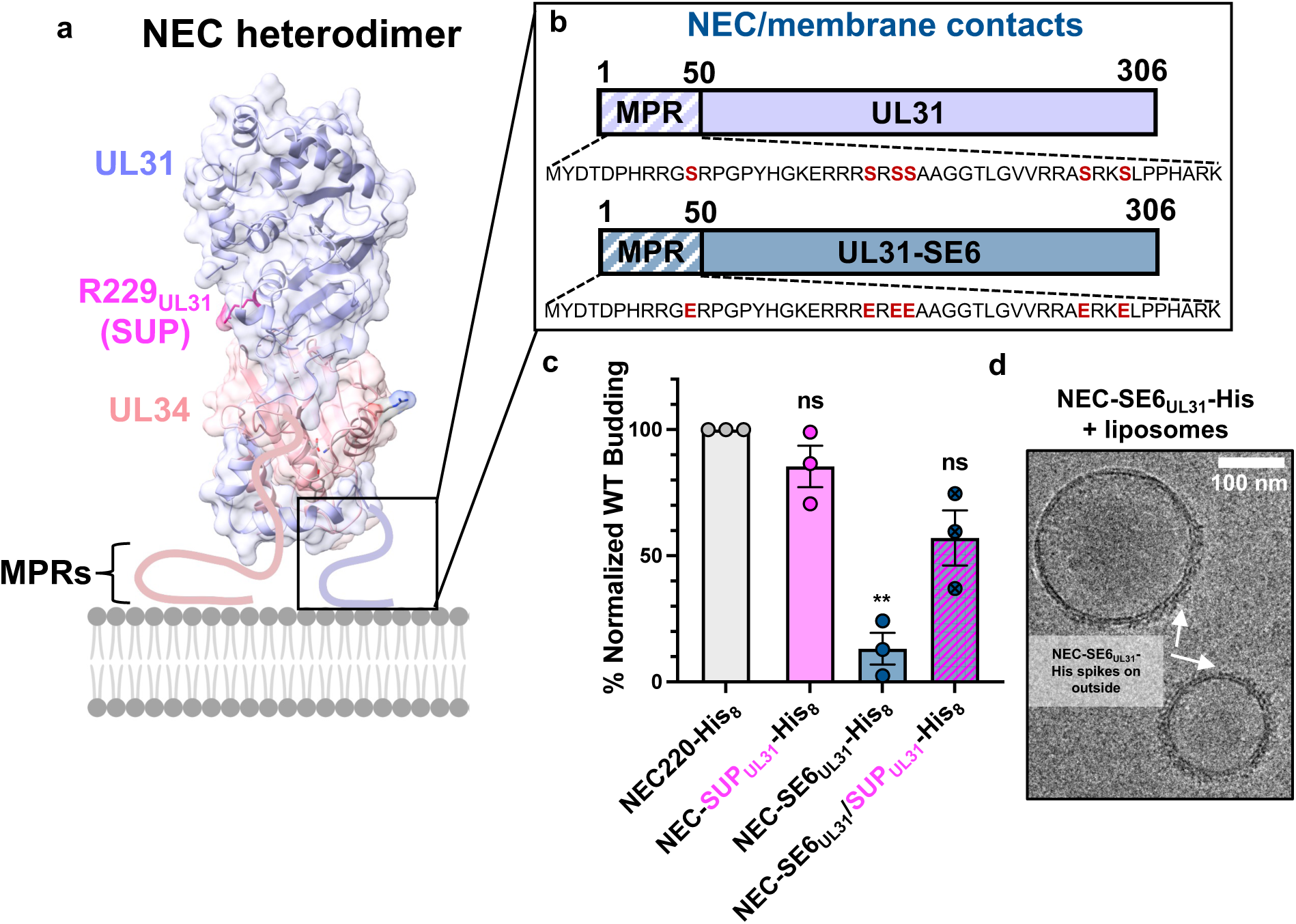
The SUP_UL31_ mutation partially restores budding to a membrane interface mutant. **a-b)** Location of membrane interface residues mutated for this study. **c)** NEC-SUP_UL31_ partially rescues budding in the membrane interface mutant *in vitro*. The percentage of *in-vitro* budding was determined by counting the number of ILVs within the GUVs after the addition of NEC220-His or the corresponding NEC mutant. A background count, the number of ILVs in the absence of NEC, was subtracted from each condition. Each construct was tested in three biological replicates, each consisting of three technical replicates. Symbols show the average budding efficiency of three technical replicates compared to NEC220-His_8_ (100%; grey). The NEC-SE6_UL31_-His_8_ data were previously reported in ^27^. Significance was calculated using an unpaired Student’s t-test with Welch’s correction (P < 0.01 = **; ns = not significant) in GraphPad Prism 9.0. **d)** Cryo-EM of NEC-SE6_UL31_-His_8_ and large unilamellar vesicles (LUVs) shows that the SE6_UL31_ mutations perturb NEC oligomerization when bound to membranes.

The NEC220 construct typically used in the *in-vitro* budding assays is soluble and depends on functional MPRs for membrane recruitment. Since the SE6_UL31_ mutation reduces NEC/membrane interactions, to bypass the defect in membrane recruitment, we used the NEC220 variant construct that contains a His_8_-tag at the C terminus of UL34 ^17,27^. When used in conjunction with membranes containing Ni-chelating lipids, the His_8_-tag ensures that the NEC220-His_8_ is recruited to the membranes even when the MPR mutations preclude membrane association. The *in-vitro* budding efficiency of NEC220-His_8_ was similar to that of untagged NEC220, suggesting that the C-terminal His_8_-tag has no deleterious effect on the membrane budding activity ^17,27^. Previously, we showed that such artificial tethering does not override the requirement for the MPR/membrane interactions and does not restore budding to MPR mutants with budding defects ^27^.

By itself, the SUP_UL31_ mutation did not change the budding efficiency of NEC220-His_8_ **(Fig. 4c)**, similar to the untagged NEC220 **(Fig. 1e).** As previously reported by our group, SE6_UL31_ mutation reduced *in-vitro* budding to ∼10% of the WT NEC220-His_8_ despite the ability to interact with membranes due to the His_8_-tag ^27^ **(Fig. 4c)**. Therefore, we performed a cryo-EM analysis to examine NEC220-SE6_UL31_-His_8_ membrane interactions. NEC220-SE6_UL31_-His_8_ was incubated with large unilamellar vesicles (LUVs) of similar composition to the GUVs used for the budding assay and imaged with cryo-EM **(Fig. 4d)**. NEC220-SE6_UL31_-His_8_ formed membrane-bound spikes on the outside of the LUVs (**Fig. 4d)**, but the internal protein coats indicative of budding ^17,19,28^ were rarely observed. This is reminiscent of the behavior of the oligomerization-deficient NEC-DN_UL34_ mutant previously reported by our group ^17^. We conclude that the SE6_UL31_ mutations perturb NEC oligomerization, likely as the consequence of weakened MPR/membrane interactions ^27^. Surprisingly, the SUP_UL31_ mutation restored budding of the SE6_UL31_ mutant to ∼50% of the WT NEC220-His_8_ **(Fig. 4c).** Therefore, the SUP_UL31_ mutation can rescue budding *in vitro,* even if partially, to a mutant that indirectly disrupts oligomerization by weakening MPR/membrane interactions.

### The SUP_UL31_ mutation does not cause major conformational changes in the NEC heterodimer

To identify the mechanism by which the SUP_UL31_ mutation can restore budding to a broad range of mutants, we first asked whether the SUP_UL31_ mutation influenced the NEC structure. We crystallized the equivalent of the previously crystallized WT NEC185Δ50 construct (UL31: 51-306 and UL34: 15-185) ^22^, which lacks the MPRs that impede crystallization and contains the SUP_UL31_ mutation, NEC185Δ50-SUP_UL31_. The NEC185Δ50-SUP_UL31_ structure was determined using molecular replacement with the WT NEC185Δ50 structure as a search model and refined to 3.9-Å resolution **(Supplementary Table S1).** NEC185Δ50-SUP_UL31_ took the space group C2_1_ with six NEC heterodimers in the asymmetric unit **(Supplementary Table S1).** The atomic coordinates and structure factors of the NEC185Δ50-SUP_UL31_ structure have been deposited to the RCSB Protein Data Bank under the accession number 8G6D.

The six non-crystallographic NEC185Δ50-SUP_UL31_ heterodimers, SUP_AB_, SUP_CD_, SUP_EF_, SUP_GH_, SUP_IJ_, and SUP_KL_ (UL34 chains: A, C, E, G, I, and K; UL31 chains: B, D, F, H, J, and were well resolved (95-99% of all residues; **Supplementary Table S2**) and structurally similar, with root mean square deviations (RMSDs) ranging from 0.67 to 1.00 Å **(Supplementary Table S3).** By contrast, the equivalent WT NEC185Δ50 construct took the P6 space group with two crystallographically independent heterodimers in the asymmetric unit, NEC_AB_ and NEC_CD_ (UL34 chains: A and C; UL31 chains: B and D) ^22^. The overall structures of the two WT NEC heterodimers and the six NEC-SUP_UL31_ mutant heterodimers are very similar. They can be superimposed with RMSDs ranging from 0.83 to 1.02 Å **(Supplementary Table S4)** and share similar heterodimeric interfaces **(Supplementary Table S5).** In four of the six SUP mutant heterodimers (SUP_AB_, SUP_CD_, SUP_EF_, and SUP_KL_), residues 129-133 and 261-268 of UL31, unresolved in the WT structure, were resolved **(Fig. S4).** All six copies of UL34 in the NEC-SUP_UL31_ mutant contained additional density at the C terminus **(Fig. S4).** Importantly, the location of residue at position 229, R229_UL31_ in WT and L229_31_ in SUP_31_, is unchanged **(Fig. S4).** Thus, the SUP_UL31_ mutation does not alter the NEC heterodimer structure in any major way.

To rule out the possibility that the SUP_UL31_ mutation altered the NEC structure only in the presence of mutations causing budding defects, we also crystallized the NEC185Δ50-DN_UL34_/SUP_UL31_ construct. The NEC185Δ50-DN_UL34_/SUP_UL31_ also took the C2_1_ space group with six heterodimers in the asymmetric unit but diffracted x-rays only to ∼6 Å resolution. Given the resolution, we did not perform an in-depth analysis on the NEC185Δ50-DN_UL34_/SUP_UL31_ mutant heterodimers within the crystals. Nonetheless, the similarities between the two constructs (the formation of hexagonal crystals, the space group, and the number of heterodimers in the asymmetric unit) suggest the DN_UL34_ mutations do not have a substantial effect on NEC crystal packing and most likely do not alter NEC conformation.

### WT NEC, NEC-SUP_UL31_, and NEC-DN_UL34_/SUP_UL31_ form similar hexagonal arrays in vitro

The WT NEC homologs from HSV-1, PRV, and HCMV oligomerize into hexagonal arrays ^17,20–23^. HSV-1 NEC220 forms membrane-bound hexagonal coats on the inner surface of budded vesicles *in vitro* ^17,28^ whereas NEC185Δ50 forms flat hexagonal lattices of very similar dimensions in the crystals ^22^. In the crystals of WT HSV-1 NEC185Δ50, the two independent NEC heterodimers, NEC_AB_ and NEC_CD_, form two very similar hexamers, hex_AB_ and hex_CD_, that are perfectly symmetrical due to the P6 crystal symmetry **(Fig. 5ab)** and have similar hexameric interfaces **(Supplementary Table S6).** However, these two hexamers form two distinct hexagonal lattices **(Fig. 5ab)** ^22^. In both lattices, interactions between the hexamers result in trimers formed by UL31/UL31 interactions **(Fig. 5ab, red).** The hex_AB_ lattice also has two types of dimers formed by either UL31/UL31 or UL31/UL34 interactions **(Fig. 5a, coral and gold,** respectively). But the hex_CD_ lattice has only one dimer type formed by UL31/UL34 interactions **(Fig. 5b, gold).** The WT HSV-1 NEC hexamers can, thus, interact in more than one way.

**Fig. 5.**
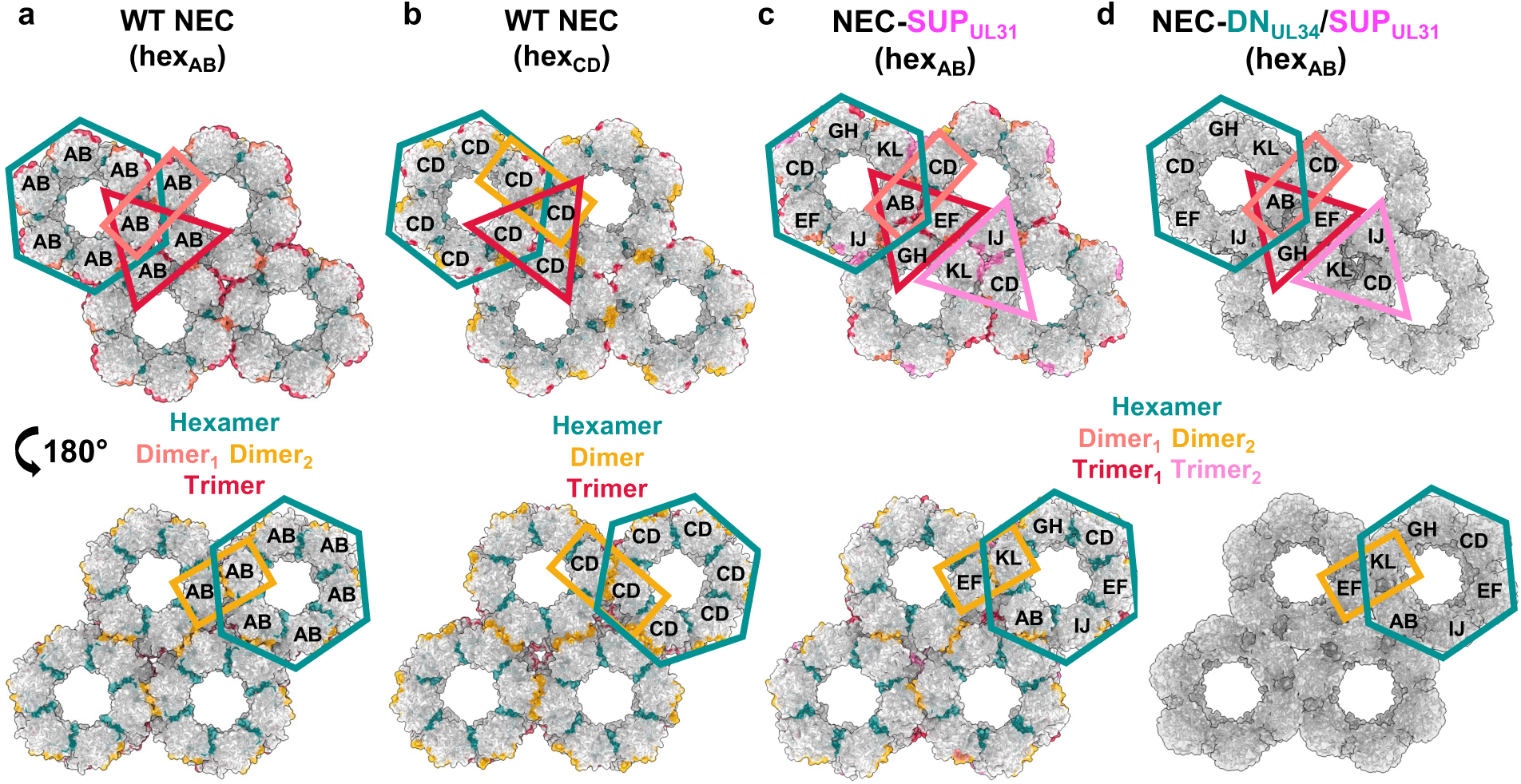
NEC-SUP_UL31_ and NEC-DN_UL34_/SUP_UL31_ form hex_AB_ lattices in the crystals. **a)** The HSV-1 WT NEC hex_AB_ (PDB ID: 4ZXS), **b)** the WT NEC hex_CD_ (PDB ID: 4ZXS), **c)** the NEC-SUP_UL31_ and **d)** the NEC-DN_UL34_/SUP_UL31_ crystal lattices ^22^. Hexameric (teal) and interhexameric (dimer_1_: coral, dimer_2_: gold, trimer_1_: red, and trimer_2_: light pink) interfaces are colored accordingly. Two distinct trimers formed in the NEC-SUP_UL31_ lattice (red and light pink). Due to resolution, an interface analysis was not performed on the NEC-DN_UL34_/SUP_UL31_ crystal lattice. The corresponding heterodimers within the lattice are labeled.

To understand how the SUP_UL31_ mutation may suppress budding defects, we examined the oligomeric arrays formed by NEC-SUP_UL31_ in the crystals and membrane-bound coats. The NEC185Δ50-SUP_UL31_ also forms hexamers in the crystals, but in this case, the hexamers are asymmetrical, being formed by six independent, non-crystallographic heterodimers in the asymmetric unit **(Fig. 5C).** Nonetheless, the NEC185Δ50-SUP_UL31_ hexamers look very similar to the WT NEC185Δ50 hexamers, with similar hexameric interfaces **(Supplementary Table S6)**, 85-97% in identity **(Supplementary Table S7)**. The SUP_UL31_ mutation thus has no major effect on the hexamer structure. The crystal lattice formed by the NEC185Δ50-SUP_UL31_ hexamers **(Fig. 5c)** resembles the WT NEC hex_AB_ lattice **(Fig. 5a; Supplementary Tables S8-S10).** The NEC185Δ50-DN_34_/SUP_31_ crystal lattice also resembles the WT NEC hex_AB_ lattice **(Fig. 5d).**

To examine the effect of the SUP_UL31_ mutation on the geometry of the membrane-bound NEC coats, we performed cryo-EM/T analyses on WT NEC220 and the mutants NEC220-SUP_UL31_ and NEC220-DN_UL34_/SUP_UL31_. Each protein complex was incubated with LUVs, similar in composition previously used to visualize the WT NEC220 coats on budded vesicles ^17,19,28^. The budded vesicles formed by NEC220-SUP_UL31_ were prone to aggregation, which reduced the number of NEC220-SUP_UL31_ particles available for data processing, resulting in a lower final resolution compared to WT NEC220 and NEC220-DN_UL34_/SUP_UL31_. Sub-tomographic averaging of the 3D reconstructions of either WT NEC220 (5.9 Å), NEC220-SUP_UL31_ (13.1 Å), or NEC220-DN_UL34_/SUP_UL31_ (5.4 Å) revealed that all three constructs formed very similar hexameric lattices **(Fig. 6a-c; Supplementary Table S11)** resembling the WT NEC hex_CD_ crystal lattice **(Fig. 5b).**

**Fig. 6.**
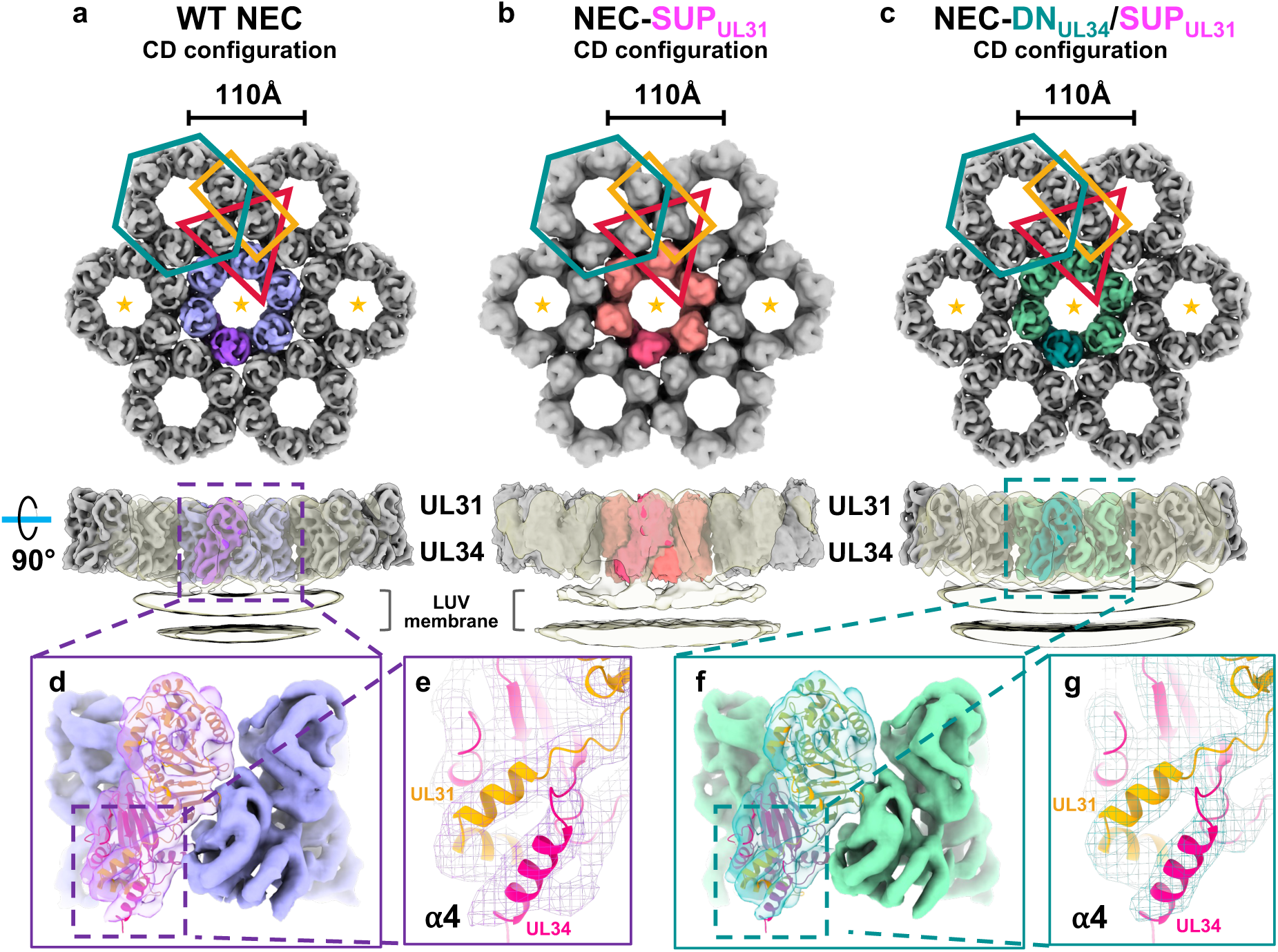
WT NEC, NEC-SUP_UL31_, and NEC-DN_UL34_/SUP_UL31_ form hex_CD_ membrane-bound coats. Cryo-ET reconstruction of **a)** the WT NEC coat at 5.9 Å, **b)** the NEC-SUP_UL31_ coat at 13.1 Å, and **c)** the NEC-DN_UL34_/SUP_UL31_ coat at 5.4 Å. Only the three hexameric units marked with orange stars are shown in lower 90°-rotated panels, where the low-pass filtered transparent densities show the connection between NEC lattices and LUV membrane. The HSV-1 NEC crystal structure (PDB ID: 4ZXS) docks similarly into both the WT NEC **(d)** and NEC-DN_UL34_/SUP_UL31_ **(f)** cryo-ET densities. **e, g)** Docking of the ⍺4 helix from the PRV NEC crystal structure (PDB ID: 4Z3U) accounts for the additional density observed in both cryo-ET reconstructions which was originally unresolved in the HSV-1 NEC crystal structure^22^.

The higher resolution of the WT NEC220 and NEC220-DN_UL34_/SUP_UL31_ averaged cryo-ET density map allowed us to dock the crystal structure of the WT NEC185Δ50 heterodimer **(Figs. 6df),** confirming that the SUP_UL31_ mutation does not perturb NEC conformation, even when bound to membranes. We also observed additional helical density at the C terminus of UL34, corresponding to helix ⍺4 that was unresolved in the WT NEC185Δ50 crystal structure ^22^ but present in the crystal structures of NEC homologs from PRV^22,31^, HCMV^23,32^, and EBV ^19^. The PRV UL34 ⍺4 helix fit well into the HSV-1 UL34 cryo-ET averages **(Figs. 6eg)**.

Interestingly, in the crystals, the WT NEC220 formed both hex_AB_ and hex_CD_ lattices whereas the mutants formed only the hex_AB_ lattice. However, in the membrane-bound coats, the WT NEC220 and both mutants formed only the hex_CD_ lattice. The reasons for these differences are yet unclear. Regardless, just as the WT NEC, both NEC-SUP_UL31_ and NEC-DN_UL34_/SUP_UL31_ mutants can form either type of lattice. Therefore, the SUP_UL31_ mutation does not promote the formation of a different NEC lattice type.

### The SUP_UL31_ mutation generates new contacts at the interhexameric interface

To determine whether the SUP_UL31_ mutation changed any contacts at the lattice interfaces, we analyzed the buried surface areas and sidechain contacts (hydrogen bonds and salt bridges) in the WT NEC and NEC-SUP_UL31_ crystal structures using PiSA interface analysis ^33^. We found that the buried surface area at 5/6 hexameric interfaces in the NEC-SUP_UL31_ crystal lattice was ∼15% smaller compared to the WT NEC hex_AB_ and hex_CD_ lattices **(Supplementary Table S12)** and that the number of hydrogen bonds and salt bridges was also reduced **(Supplementary Table S13).** The remaining hexameric interface, A/K, had more contacts than the WT **(Supplementary Table S13).** Thus, the SUP_UL31_ mutation appears to alter contacts at the hexameric interfaces.

By contrast, the buried surface area at the interhexameric interfaces was larger in the NEC-SUP_UL31_ crystal lattice **(Supplementary Table S14)** and had additional interactions absent from the WT NEC lattice **(Supplementary Table S15).** For example, the trimeric interface in the WT NEC hex_AB_ lattice has one salt bridge, E138_UL31_-R155_UL31_ **(Fig. 7a).** The NEC-SUP_UL31_ lattice has two trimeric interfaces, B/F/H and D/J/L. In both, E138_UL31_ forms a salt bridge with another residue, R193_UL31_ **(Fig. 7b).** R155_UL31_ forms a salt bridge with E267_UL31_, but only in the B/F/H trimer **(Fig. 7b).** In the D/J/L trimer, these residues are ∼7 Å apart. E267_UL31_ was unresolved in the WT NEC_AB_ structure, but its side chain is too far away from the interhexameric interface to participate in any contacts **(Fig. 7a)**. As another example, there is a new salt bridge, D286_UL31_-R295_UL31_, at the NEC-SUP_UL31_ dimeric interfaces B/D and F/L **(Fig. 7b),** which is absent from the WT NEC lattice. Therefore, the NEC-SUP_UL31_ mutation causes the formation of additional contacts at the interhexameric interface that could ostensibly stabilize the NEC lattice even in the presence of lattice-destabilizing mutations. New interhexameric contacts likely also form in the hex_CD_ lattice, but in the absence of higher resolution data, what residues participate in these contacts is unknown.

**Fig. 7.**
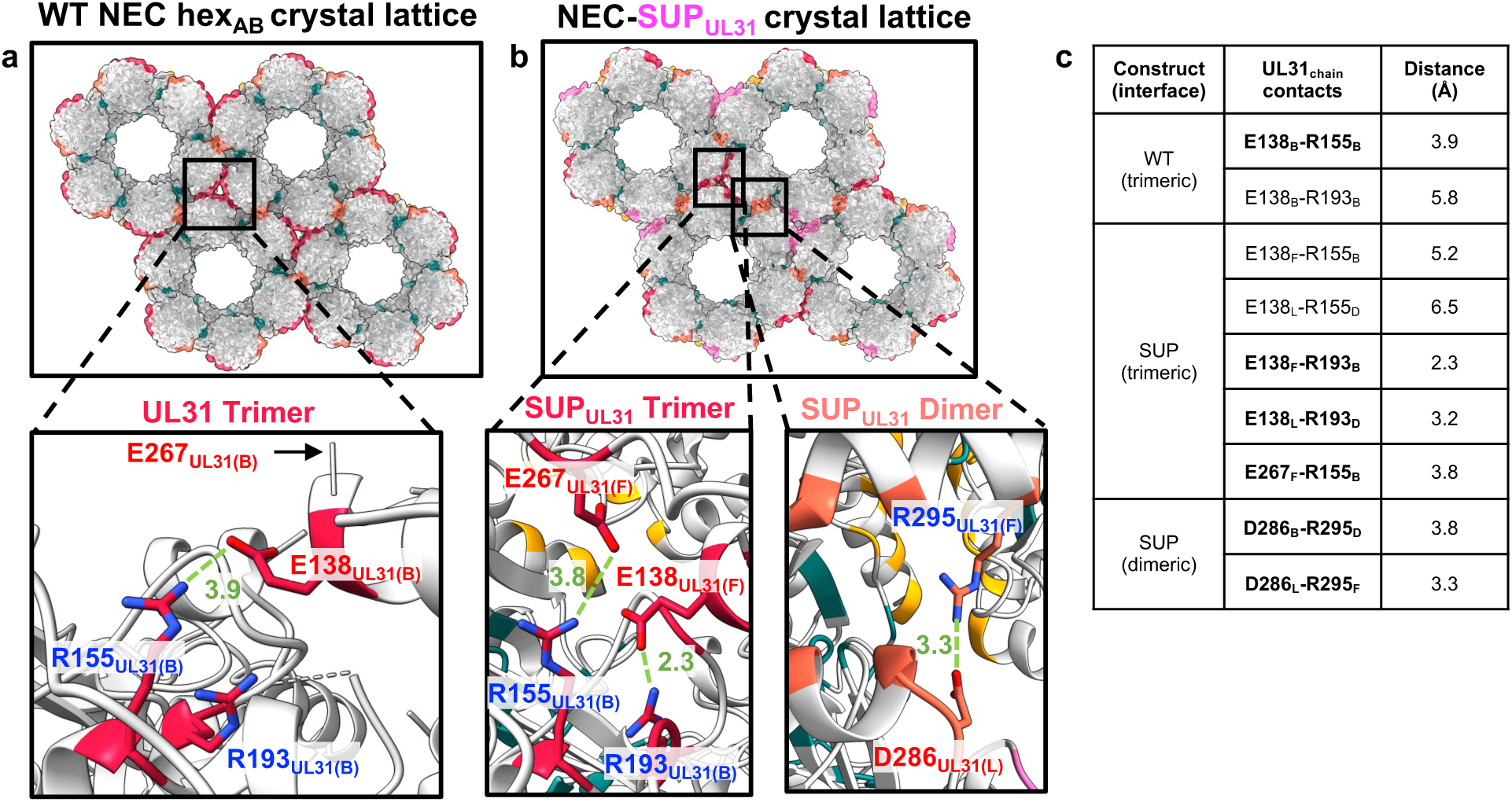
The interhexameric interfaces in WT NEC hex_AB_ and NEC-SUP_UL31_ lattices have different contacts. Close-up views of the interhexameric interfaces in the **(a)** WT NEC crystal lattice (UL31 trimer; crimson) and **(b)** NEC-SUP_UL31_ crystal lattice [UL31 trimers (crimson and pink) and dimers (orange and yellow)]. The HSV-1 NEC crystal structure (PDB: 4ZXS) was used to generate the figure in panel **a**. Salt bridges are shown as green dashed lines, with distances in Angstrom. Residues forming salt bridges are shown as sticks and colored in blue (Rs) or red (Es and Ds). **c)** Distances of contacts at the highlighted interfaces. Contacts in bold are salt bridges.

### The SUP_UL31_ mutation generates new interface contacts indirectly by eliminating a salt bridge

While residue 229_UL31_ does not participate in any interface contacts directly, it is located right above the heterodimeric globular interface **(Fig. 8a).** In the WT NEC_AB_ structure, the R229_UL31_ side chain makes a salt bridge with the nearby D129_UL31_. D129_UL31_ is located at one end of a mostly disordered loop, residues 129_UL31_-134_UL31_ **(Fig. 8ac)**, that was only partially resolved in the WT NEC_AB_ structure and unresolved in the WT NEC_CD_ structure. The R229L_UL31_ mutation eliminates the salt bridge, and in all SUP_UL31_ heterodimers (except for SUP_KL_ where it was unresolved), the D129_UL31_ side chain points away from L229_UL31_ **(Fig. 8b).** Interestingly, in the NEC-SUP_UL31_ structure, the 129_UL31_-134_UL31_ loop is better ordered. It was fully resolved in four out of six NEC-SUP_UL31_ heterodimers **(Fig. 8d, red)** and partially resolved in the remaining two **(Supplementary Table S9).** Moreover, this loop also participates in new contacts at the interhexameric interface in five out of the six NEC-SUP_UL31_ heterodimers, **(Supplementary Table S9).** To sum up, in the NEC-SUP_UL31_, the lack of the R229_UL31_-D129_UL31_ salt bridge correlates with a more ordered 129_UL31_-134_UL31_ loop **(Fig. 8bd, red)** and a larger interhexameric interface **(Fig. 8d)**. Therefore, we hypothesize that by eliminating the salt bridge, the SUP_UL31_ mutation releases the loop, which then forms new contacts at the interhexameric interface.

**Fig. 8.**
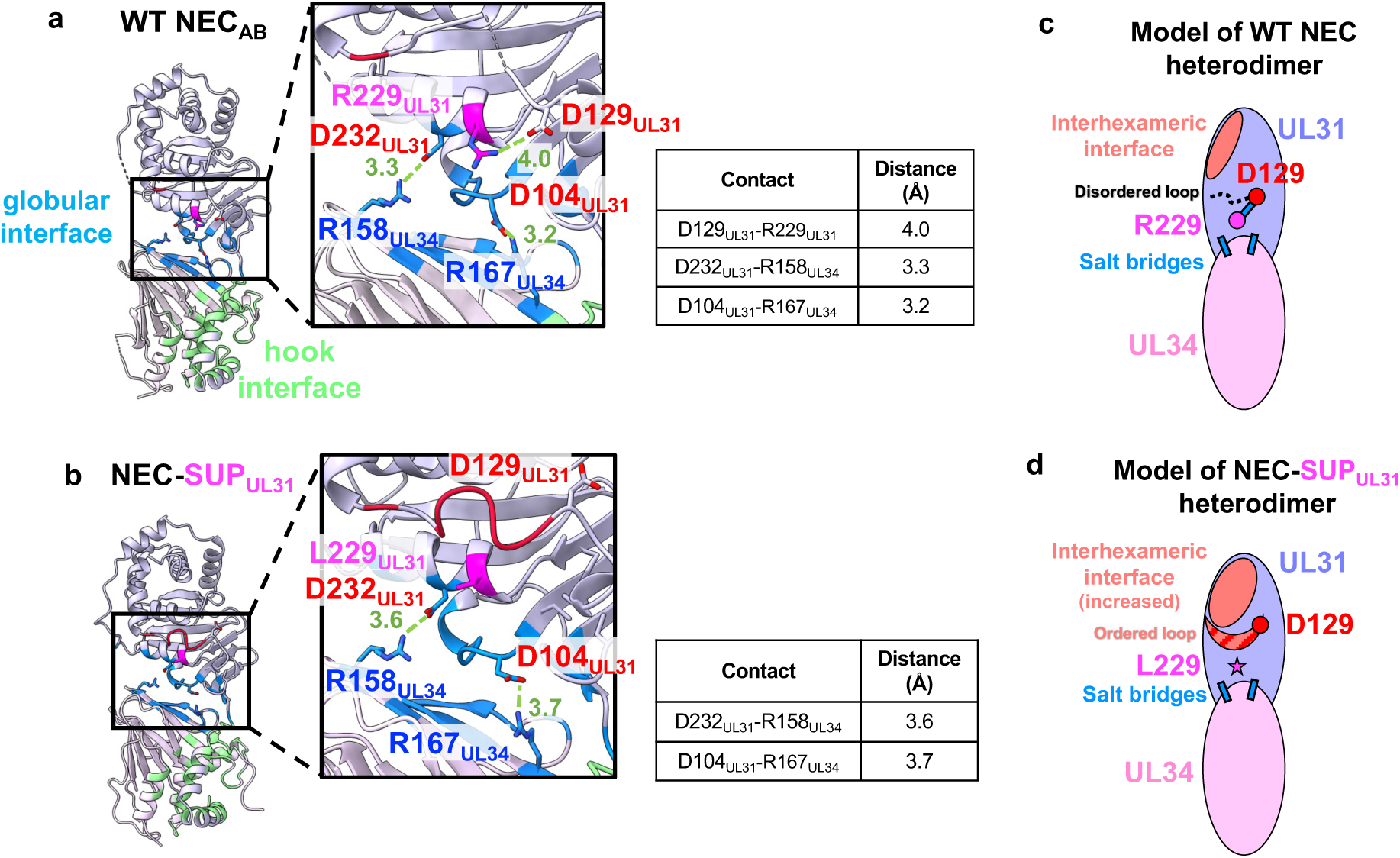
The SUP_UL31_ mutation introduces changes to the heterodimeric interface. The crystal structures of **(a)** WT NEC_AB_ and **(b)** NEC-SUP_UL31_ heterodimers. The heterodimeric interface residues are colored blue (globular interface) or green (hook interface). Residue 229_UL31_ is colored magenta. Insets show close-up views of the globular interface. Residues forming salt bridges at the interface are shown as sticks and colored in blue (Rs) or red (Es and Ds). Residue 229_UL31_ is shown as sticks and colored in magenta. Salt bridges are shown as dashed green lines, with distances in Angstrom. Distances are also listed in the corresponding tables. The resolved portions of the dynamic loop 129_UL31_-134_UL31_ are shown in red. The HSV-1 NEC crystal structure (PDB: 4ZXS) was used to generate the figure in (**a)**. Corresponding cartoon models of either the **(c)** WT NEC or **(d)** NEC-SUP_UL31_ heterodimeric interfaces.

## DISCUSSION

Herpesviruses translocate their capsids from the nucleus to the cytoplasm by an unusual mechanism that requires the formation of membrane-bound coats by the virally encoded heterodimeric complex, the NEC ^17,20,21^. The coats are composed of a hexagonal NEC lattice, and mutations that disrupt the lattice interfaces reduce budding *in vitro* ^17,22,27^ and viral replication ^24–26,30^, attesting to its central role in the NEC membrane-budding function. Here, we demonstrated that a suppressor mutation within the UL31 protein, SUP_UL31_, restored efficient membrane budding *in vitro* and viral replication to a broad range of budding-deficient NEC mutants. The SUP_UL31_ mutation thus acts as a universal suppressor of membrane budding defects in NEC. Furthermore, we found that the SUP_UL31_ mutation expanded lattice interfaces by indirectly creating new interface contacts. We hypothesize that the SUP_UL31_ mutation exerts its powerful suppressor effect by stabilizing the hexagonal coats destabilized by mutations.

### The SUP_UL31_ mutation promotes the formation of new interhexameric contacts

Since the SUP_UL31_ mutation rescued budding defects caused by disruptions of the hexagonal lattice, we initially hypothesized that it may do so promoting the formation of a different, potentially, non-hexagonal lattice with distinct interfaces. Instead, we found that NEC-SUP_UL31_ and NEC-DN_UL34_/SUP_UL31_ mutants formed hexagonal lattices both in the crystals and in membrane-bound coats that were very similar to those formed by the WT NEC. However, the interhexameric interfaces in the NEC-SUP_UL31_ crystal lattice are larger due to several new interactions, particularly, salt bridges. The SUP_UL31_ mutation does not participate in any interface contacts itself. Instead, we hypothesize that it promotes new interface contacts indirectly by eliminating the R229_UL31_-D129_UL31_ salt bridge **(Fig. 8b)**. This releases the 129_UL31_-134_UL31_ loop, which becomes better ordered **(Fig. 8d)** and forms new contacts at the interhexameric interface **(Fig. 7b).** Larger interhexameric lattice interfaces would be expected to reinforce the lattice. By stabilizing the lattice disrupted by mutations, the SUP_UL31_ mutation could restore efficient budding.

It is easy to envision how the SUP_UL31_ mutation might suppress budding defects caused by mutations that destabilize interhexameric interactions by compensating for the loss of those interactions locally with new interhexameric interactions. However, the SUP_UL31_ mutation also suppresses budding defects of mutants that destabilize the hexamers themselves. Therefore, we propose a more general mechanism for this type of suppression. We suggest that the NEC hexamers weakened by interface mutations can be stabilized not only by the interactions between adjacent NEC heterodimers within the hexamer itself but also by their incorporation into a larger lattice where interhexameric contacts would limit the dissociation of NEC heterodimers from the hexamer (**Fig. 9**). By strengthening these latter contacts, the SUP_UL31_ mutation can thereby compensate for different kinds of lattice defects.

**Fig. 9.**
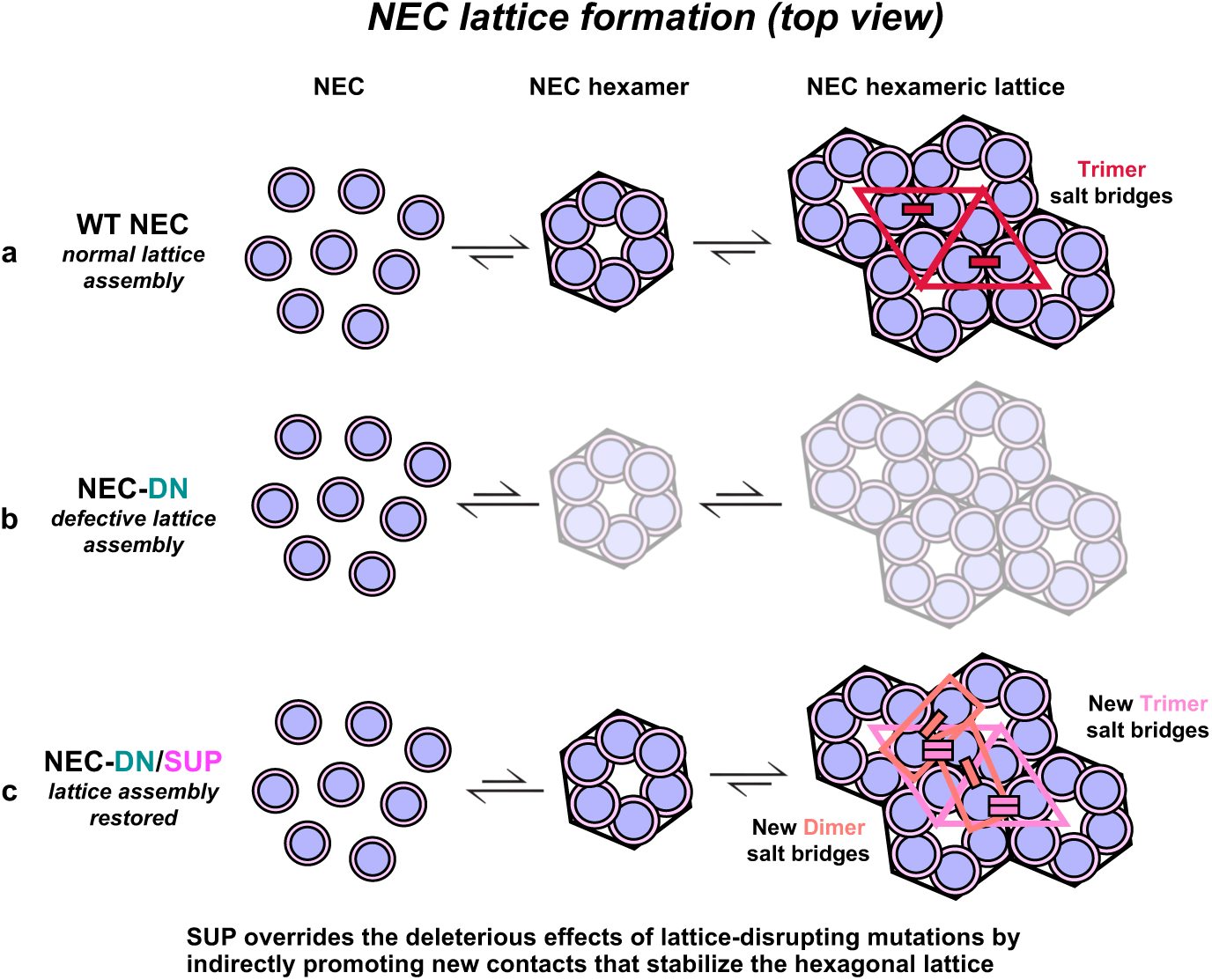
A model of SUP_UL31_ budding restoration in the context of a lattice destabilizing mutation. **a)** WT NEC heterodimers arrange into hexamers that build into a stable hexameric lattice by forming contacts at the hexameric and interhexameric interfaces. The salt bridges located at the interhexameric interface trimers (crimson) favor lattice association, rather than disassociation. In the presence of a lattice destabilizing mutation such as NEC-DN_UL34_ **(b)**, hexamer formation and lattice assembly are perturbed. In contrast, the SUP_UL31_ mutation **(c)** restores lattice formation by promoting the formation of new contacts at both the dimeric and trimeric interhexameric interfaces (pink and salmon), resulting in a stable and functional NEC lattice, despite the presence of a destabilizing mutation.

### The SUP_UL31_ mutation makes the NEC more conformationally dynamic

The WT NEC185Δ50 crystallized in the P6 space group, in which the hexamers are perfect, being related by crystallographic symmetry. However, both the NEC185Δ50-SUP_UL31_ and NEC185Δ50-DN_UL34_/SUP_UL31_ mutants crystallized in the C2_1_ space group, in which the hexamers are non-crystallographic and, thus, imperfect. This suggested that the NEC heterodimer becomes more conformationally dynamic in the presence of the SUP_UL31_ mutation. Indeed, the heterodimeric UL31/UL34 interface between the globular cores of UL31 and UL34 buries a ∼6-20% larger surface area in all six NEC-SUP_UL31_ heterodimers compared to the WT NEC heterodimers **(Fig. S4, blue; Supplementary Table S16),** suggesting flexibility at this interface.

Residue 229_UL31_ is located between two intermolecular salt bridges at the heterodimeric interface formed by the UL31 and UL34 globular cores, R158_UL34_-D232_UL31_ and R167_UL34_-D104_UL31_ **(Fig. 8b).** Previous molecular dynamics (MD) study proposed that these salt bridges contribute to the overall stability of the HSV-1 NEC heterodimer ^34^. The HCMV NEC, which has only one salt bridge, was found to be more dynamic than HSV-1 in MD simulations^34^. The EBV NEC, which also has only one salt bridge, is a conformationally dynamic heterodimer as revealed by its crystal structure ^19^. The number of intermolecular salt bridges at the globular UL31/UL34 interface thus correlates with the stability of the NEC heterodimer.

We propose that the intramolecular salt bridge R229_UL31_-D129_UL31_ is another important contributor to the stability of the NEC heterodimer. Although the NEC-SUP_UL31_ still has two intermolecular salt bridges at the heterodimeric interface, it lacks the intramolecular salt bridge **(Fig. 8b)**, which increases its flexibility. This increased flexibility could explain why both NEC-SUP_UL31_ and NEC-DN_UL34_/SUP_UL31_ required extra additives and small molecules (from the Silver Bullets) to form crystals relative to the WT NEC ^22^ and why both NEC-SUP_UL31_ and NEC-DN_UL34_/SUP_UL31_ took the lower symmetry space group C2_1_, rather than P6.

Despite the proposed variations in flexibility among closely related NECs, only two types of lattice configurations have been observed, hex_AB_ and hex_CD_. HSV-1 NEC, be it the WT or the two mutants presented in this study, formed either hex_AB_ (WT-NEC, NEC-SUP_UL31_, and NEC-DN_UL34_/SUP_UL31_) or hex_CD_ (WT-NEC) lattice configurations in crystals and only hex_CD_ lattice in membrane-bound coats. HCMV NEC packed into a hexagonal lattice resembling the hex_CD_ lattice configuration ^23^. Conversely, PRV NEC formed a hexagonal lattice resembling the hex_AB_ lattice configuration in membrane-bound coats ^20^. HCMV and PRV NEC homologs could, in principle, adopt alternative configurations, hex_AB_ or hex_CD_, respectively, under different experimental conditions. Regardless, how the NEC assembles into a hexameric lattice of either configuration is still unknown. The two different lattice types could be assembled via different routes, yet their biological relevance is still unclear.

### The SUP_UL31_ mutation acts as a universal suppressor against mutations disrupting the NEC budding activity

The SUP_UL31_ mutation was initially identified as an extragenic suppressor of a nuclear budding defect caused by a double mutation within UL34, D35A_UL34_/E37A_UL34_ (DN_UL34_), in HSV-1 infected cells ^25^. Subsequently, we showed that the DN_UL34_ mutation blocks the formation of the hexagonal NEC lattice ^17^ by eliminating important polar contacts at the hexameric interface ^22^ and that the SUP_UL31_ mutation restored effective membrane budding *in vitro* to the DN_UL34_ and another hexameric interface mutant, V92F_UL34_ ^22^. Here, we found that the SUP_UL31_ mutation could restore budding to many lattice interface mutants (T123Q_UL34_, F252Y_UL31,_ and E153R_UL31_; **Fig. 1**); the heterodimeric interface mutants (K137A_UL34_ and K137A/R139A_UL34_; **Fig. 3**); and even to a membrane interface mutant (SE6_UL31_; **Fig. 4**). Thus, the SUP_UL31_ mutation acts a universal suppressor mutation that restores efficient budding *in vitro* and, in several cases, viral replication to a diverse range of budding-deficient NEC mutants.

Although mutations that cause budding defects target diverse interfaces, all are expected to destabilize the hexagonal NEC lattice, which is essential for the membrane budding process. The destabilizing effect of the lattice interface mutations is the most apparent. But mutations destabilizing the NEC heterodimer would also be expected to weaken the lattice by destabilizing its core building block. Finally, NEC/membrane interactions likely destabilize the lattice within the membrane-bound NEC coat indirectly. By reinforcing the lattice, the SUP_UL31_ mutation could overcome these lattice destabilizing defects regardless of their nature.

In some cases, the SUP_UL31_ mutation could fully restore the *in-vitro* budding activity but not viral replication, e.g., in the presence of the T123Q_UL34_ and E153R_UL31_ **(Fig. 2)**. We hypothesize that these mutations may perturb the NEC functions that do not involve membrane deformation, e.g., nuclear lamina dissolution, capsid docking at the INM, or capsid recruitment. Thus, the SUP_UL31_ mutation specifically restores budding defects.

If the SUP_UL31_ mutation forms a stronger hexagonal lattice, why isn’t this mutation positively selected? We hypothesize that this mutation may impair other functions of the NEC. This idea is supported by the observation that in the *in-trans* complementation experiment, viral replication in the presence of the SUP_UL31_ mutation does not reach the WT levels **(Fig. 2).** While the SUP_UL31_ mutation is not positively selected, it provides the virus with a strategy to maintain replication in the presence of external stressors such as inhibitors targeting the NEC. Further work to identify and characterize other herpesviral suppression mechanisms could aid in the advancement of novel herpesviral therapeutics.

## MATERIALS AND METHODS

### Cells and viruses

Vero and Hep-2 cells were maintained as previously described ^9^. The properties of HSV-1(F) and vRR1072(TK+) (a UL34-null virus derived by homologous recombination with HSV-1(F) have also been previously described ^9,35^. The UL34-null virus and the UL34-null/SUP_UL31_ recombinant viruses used for complementation assays were derived from the pYEbac102 clone of the HSV-1 strain (F) genome in the bacterial strain GS1783 (a gift from G. Smith, Northwestern U.) ^36–38^ as previously described. All UL34-null viruses were propagated on Vero tUL34 CX cells that express HSV-1 pUL34 under the control of its native promoter regulatory sequences ^39^. Vero tUL34 CX cells were propagated in DMEM high glucose supplemented with 5% fetal bovine serum and the antibiotic penicillin and streptomycin.

### Cloning

All primers used for cloning are listed in **Supplementary Table S17**. Cloning of UL31 (1-306), UL34 (1-220), UL34 (15-185), UL34-His_8_ (1-220 with a C-terminal His_8_-tag) and the corresponding UL31 and UL34 mutants [R229L_UL31_ (SUP_UL31_), D35A_UL34_/E37A_UL34_ (DN_UL34_), E37A_UL34_, T123Q_UL34_, F252Y_UL31_, and E153R_UL31_, and S11E_UL31_/S24E_UL31_/S26E_UL31_/S27E _UL31_/S40E_UL31_/S43E_UL31_ (SE6_UL31_)] was previously described ^17,22,27^.

#### Oligomeric interface mutants

Site-directed mutagenesis of pJB14 (UL31 1-306 R229L_UL31_) was performed using splicing-by-overlap extension protocol followed by restriction digest into the pKH90 plasmid (containing an N-terminal His-SUMO-PreScission tag in-frame with a BamHI restriction site) to create either the F252Y_UL31_/R229L_UL31_ (pED20) or E153R_UL31_/R229L_UL31_ (pED21) double mutants.

#### Heterodimeric interface mutants

Site-directed mutagenesis of pJB02 (UL34 1-220) was performed using splicing-by-overlap-extension protocol followed by restriction digest into the pJB02 plasmid (containing an N-terminal GST-PreScission tag in-frame with a SalI restriction site) to create either the K137A_UL34_ (pED25), R139A_UL34_ (pED26), or the K137A_UL34_/R139A_UL34_ (pED27) mutants.

#### Membrane interface mutants

Site-directed mutagenesis of pJB60 (UL31-SE6_UL31_ 1-306) was performed using an inverse PCR protocol followed by blunt-end ligation to create the SE6_UL31_/R229L_UL31_ mutant (pED45).

#### Crystallization constructs

Digested PCR fragments containing R229L_UL31_Δ50-306 were amplified from pJB114 (UL31 1-306 R229L_UL31_) and subcloned by restriction digest into a pET24b(+) plasmid harboring an N-terminal His_6_-SUMO-PreScission tag in-frame with a BamHI restriction site plasmid to create the R229L_UL31_Δ50-306 plasmid (pXG20). Digested PCR fragments containing D35A_UL34_/E37A_UL34_ were amplified from pJB06 (UL34 1-246 D35A_UL34_/E37A_UL34_) and subcloned by restriction digest into a pGEX-6P1 vector containing an N-terminal GST-PreScission tag in-frame with a SalI restriction site to create the UL34 15-185 D35A_UL34_/E37A_UL34_ plasmid (pJB66).

#### Cell complementation constructs

Plasmid pRR1072Rep, which was the parent vector for UL34 mutant plasmids used for cell culture complementation assays has been previously described ^26^. Mutant derivatives of pRR1072Rep that carry the D35A_UL34_/E37A_UL34_ and K137A_UL34_/R139A_UL34_ double mutations have also been previously described and were referred to as CL04 and CL10 in that publication ^26^. Derivatives of pRR1072Rep containing the single D35A_UL34_, E37A_UL34_, T123Q_UL34_, K137A_UL34,_ and R139A_UL34_ were constructed by Gibson assembly. Plasmids were assembled from two PCR products, each generated using pRR1072Rep as a template and using either a mutagenic forward primer paired with a reverse primer from the ampicillin resistance gene, or a mutagenic reverse primer paired with a forward primer from the ampicillin resistance genes. PCR products were digested with DpnI to remove template sequences and then assembled using the New England BioLabs 2X Gibson assembly master mix according to the manufacturer’s instructions.

### Expression and purification of WT NEC, oligomeric interface, and heterodimeric interface mutants

Plasmids encoding HSV-1 UL31 1-306 (pKH90) and UL34 1-220 were co-transformed into *Escherichia coli* BL21(DE3) LoBSTr cells (Kerafast) to generate wild-type NEC220 ^17,28^. All mutant constructs contained UL31 1-306 and UL34 1-220 amino acid boundaries. Plasmids encoding the appropriate mutations of either UL31 or UL34 were also co-transformed into *E. coli* BL21(DE3) LOBSTR cells (Kerafast) to generate the various NEC oligomeric and heterodimeric interface mutants (listed in **Supplementary Table S18**). The expression and purification of NEC220 and some oligomeric interface mutants (NEC-DN_UL34_, NEC-DN_UL34_/SUP_UL31_, NEC-SUP_UL31_, NEC-E37A_UL34_, NEC-T123Q_UL34_, NEC-F252Y_UL31_, and NEC-E153R_UL31_) were described previously ^17,22^. The expression and purification of oligomeric interface mutants (NEC-SUP_UL31_/F252Y_UL31_, NEC-SUP_UL31_/E153R_UL31_, and NEC-SUP_UL31_/T123Q_UL34_) and heterodimeric interface mutants (NEC-K137A_UL34_, NEC-R139A_UL34_, NEC-K137A_UL34_/R139A_UL34,_ NEC-K137A_UL34_/SUP_UL31_ and NEC-K137A_UL34_/R139A_UL34_/SUP_UL31_) are described below. Cells expressing the corresponding NEC construct were expressed using auto-induction at 37 °C in TB supplemented with 100 μg/mL ampicillin, 100 μg/mL kanamycin, and 34 μg/mL chloramphenicol, 0.2% lactose, and 2 mM MgSO_4_ for 4 h. The temperature was reduced to 25 °C for 16 h. Cells were harvested at 5,000 x g for 30 min.

All purification steps were performed at 4 °C, as previously described ^17,28^. Cell pellets were resuspended in lysis buffer (50 mM Na HEPES pH 7.5, 500 mM NaCl, 1 mM TCEP, and 10% glycerol) supplemented with Complete protease inhibitor (Roche) and lysed using a microfluidizer (Microfluidics). The cell lysate was clarified by centrifugation at 18,000 x g for 40 min and passed over a Ni Sepharose 6 column (Cytiva) equilibrated with lysis buffer. The protein-bound column was washed with 20 mM and 40 mM imidazole lysis buffer and bound proteins were eluted with 250 mM imidazole lysis buffer. Eluted proteins were passed over a Glutathione Sepharose 4B column and washed with lysis buffer. The His_6_-SUMO and GST tags were cleaved for 16 h by PreScission Protease, produced in-house from a GST-PreScission fusion expression plasmid (a gift of Peter Cherepanov, Francis Crick Institute). The protein was passed over 2 x 1 mL HiTrap Talon columns (Cytiva) to remove His_6_-SUMO, followed by injection onto a size-exclusion column (Superdex 75 10/300; Cytiva) equilibrated into gel-filtration buffer (20 mM Na HEPES, pH 7.0, 100 mM NaCl, and 1 mM TCEP), as previously described. Fractions containing pure protein, as assessed by 12% SDS-PAGE and Coomassie staining were pooled and concentrated as described below.

For both NEC-SUP_UL31_ and NEC-DN_UL34_/SUP_UL31_, the cleaved proteins were passed over a HiTrap SP XL (5 mL; Cytiva) ion-exchange column, to remove free His_6_-SUMO. Bound proteins were eluted using a linear salt gradient (60 mL) made from no salt gel filtration buffer (20 mM Na HEPES, pH 7.0, and 1 mM TCEP) and salt gel filtration buffer (20 mM Na HEPES, pH 7.0, 1 M NaCl, and 1 mM TCEP). Proteins typically eluted ∼ 360 mM NaCl, at which point the gradient was held constant until the UV signal returned to baseline. Fractions containing pure protein, as assessed by 12% SDS-PAGE and Coomassie staining, were pooled and diluted using no salt gel filtration buffer to reach a 100 mM NaCl concentration, which is required for downstream liposome budding experiments described below. For all purifications described herein, the protein was concentrated to ∼ 1 mg/mL and stored at −80 °C to prevent degradation observed at 4 °C. Protein concentrations were determined by the absorbance at 280 nm. A typical yield was ∼0.5 mg/L.

### Expression and purification of membrane interface constructs

Plasmids containing UL31 1-306 (pKH90) and UL34 1-220-His_8_ (pJB57) were co-transformed into *E. coli* BL21(DE3) LoBSTr cells (Kerafast) to generate NEC220-His_8_. All the following constructs contained UL31 1-306 and UL34 1-220 amino acid boundaries. A list of plasmids co-transformed to create the NEC-SE6_UL31_-His_8_, NEC-SUP_UL31_-His_8_, and NEC-SE6_UL31_/SUP_UL31_-His_8_ constructs are listed in **Supplementary Table 18**. Cells expressing the corresponding NEC mutant were grown using auto-induction at 37 °C in TB supplemented with 100 μg/mL ampicillin, 100 μg/mL kanamycin, and 34 μg/mL chloramphenicol, 0.2% lactose, and 2 mM MgSO_4_ for 4 h. The temperature was reduced to 25 °C for 16 h. Cells were harvested at 5,000 x g for 30 min.

Cells were resuspended in lysis buffer supplemented with Complete protease inhibitor (Roche) and lysed using a microfluidizer (Microfluidizer). The cell lysate was clarified by centrifugation at 18,000 x g for 40 min and passed over a Ni Sepharose 6 column (Cytiva). The column was washed with 20 mM and 40 mM imidazole lysis buffer and bound proteins were eluted with 250 mM imidazole lysis buffer. Eluted proteins were passed over a Glutathione Sepharose 4B column and washed with lysis buffer. The His_6_-SUMO and GST tags were cleaved for 16 h by PreScission Protease, produced in-house from a GST-PreScission fusion expression plasmid. The protein was loaded on a size-exclusion column (Superdex 75 10/300; Cytiva) equilibrated with a gel-filtration buffer. Fractions containing pure protein, as assessed by 12% SDS-PAGE and Coomassie staining were pooled and concentrated as described above.

### Expression and purification of crystallization constructs

SUP_UL31_Δ50-306 (pXG20) and UL34 15-185 (pJB04) or SUP_UL31_Δ50-306 (pXG20) and UL34 15-185 D35A_UL34_/E37A_UL34_ (pJB66) plasmids were co-transformed into *E. coli* BL21(DE3) LoBSTr cells (Kerafast) to produce the NEC-SUP_UL31_ and NEC-DN_UL34_/SUP_UL31_ crystallization constructs, respectively. Cells expressing the corresponding NEC mutant were grown using auto-induction at 37 °C in TB supplemented with 100 μg/mL ampicillin, 100 μg/mL kanamycin, and 34 μg/mL chloramphenicol, 0.2% lactose, and 2 mM MgSO_4_ for 4 h. The temperature was reduced to 25 °C for 16 h. Cells were harvested at 5,000 x g for 30 min.

Cells were resuspended in lysis buffer supplemented with Complete protease inhibitor (Roche) and lysed using a microfluidizer (Microfluidizer). The cell lysate was clarified by centrifugation at 18,000 x g for 40 min and passed over a Ni Sepharose 6 column (Cytiva). The column was washed with 20 mM and 40 mM imidazole lysis buffer and bound proteins were eluted with 250 mM imidazole lysis buffer. Eluted proteins were passed over a Glutathione Sepharose 4B column and washed with lysis buffer. The His_6_-SUMO and GST tags were cleaved for 16 h by PreScission Protease, produced in-house from a GST-PreScission fusion expression plasmid. The protein was passed over 2 x 1 mL HiTrap Talon columns (Cytiva) to remove His_6_-SUMO, followed by injection onto a size-exclusion column (Superdex 75 10/300; Cytiva) equilibrated into gel-filtration buffer, as previously described. Fractions containing pure protein, as assessed by 12% SDS-PAGE and Coomassie staining were pooled and concentrated as described above.

### In-vitro budding assays

Giant unilamellar vesicles (GUVs) were prepared as previously described ^17^. For budding quantification, 10 μL of POPC:POPA:POPS=3:1:1 (Avanti Polar Lipids) GUVs containing 0.2% ATTO-594 DOPE (ATTO-TEC GmbH) fluorescent dye were added to gel filtration buffer containing 0.2 mg/mL (final concentration) Cascade Blue Hydrazide (ThermoFisher Scientific) and either 1 μM WT NEC or NEC mutant (final concentration), for a total sample volume of 100 μL. Reactions incubated for 5 min at 20 °C before imaging in 96-well chambered coverglass (Brooks Life Science Systems). Samples were imaged using a Nikon A1R Confocal Microscope with a 60x oil immersion lens at the Imaging and Cell Analysis Core Facility at Tufts University School of Medicine. Budding events were quantified by manually counting ∼300 vesicles total in 15 different frames of the sample. Before analysis, the background (GUVs in the absence of NEC) was subtracted from the raw values. All data values are reported in the **Source Data File**. Each sample was tested in at least three biological replicates, each containing three technical replicates. Reported values represent the average budding activity relative to NEC220 or NEC220-His_8_ (100%). The standard error of the mean is reported for each measurement. Significance was calculated using an unpaired one-tailed *t*-test against NEC220. Statistical analyses and data presentation were performed using GraphPad Prism 9.1.0.

### Complementation assays

24-well cultures of Hep-2 cells at 70% confluence were transfected with 0.05 μg of pCMVβ, expressing the β-galactosidase gene, and 0.25 μg of wild-type or mutant UL34 plasmid using Lipofectamine as described by the manufacturer (Gibco-BRL) and incubated at 37°C overnight. The cells were then infected with 10 PFU of the BAC-derived UL34-null virus or UL34-null/SUP_UL31_ virus per cell and incubated at 37°C for 90 min. Monolayers were washed once with pH 3 sodium citrate buffer (50 mM sodium citrate, 4 mM potassium chloride, adjusted to pH 3 with hydrochloric acid) and then incubated at room temperature in fresh citrate buffer for one minute. Cells were washed with V medium (Dulbecco’s modified Eagle’s medium, penicillin-streptomycin, 1% heat-inactivated calf serum) two times. One milliliter of V medium was then added to each well, and after 16 h of incubation at 37 °C, cell lysates were prepared by freezing and thawing followed by sonication for 20 s at power level 2 with a Fisher sonic dismembrator. The amount of infectivity in each lysate was determined by plaque assay titration on UL34-complementing cells. Part of each cell lysate was assayed for β-galactosidase expression as previously described ^26^. Transfection efficiencies in all samples were within 20% of each other. Each sample was tested in at least three biological replicates, each containing one technical replicate. The raw titers and log PFU values for each biological replicate are reported in the **Source Data File**.

### Evaluation of mutant protein expression in mammalian cells

12-well cultures of Hep-2 cells at 70% confluence were transfected with 0.5 μg of wild-type or mutant UL34 or UL31-FLAG plasmid using Lipofectamine as described by the manufacturer (Gibco-BRL) and incubated at 37 °C overnight. The cells were then infected with 10 PFU of the corresponding null virus and incubated at 37 °C for 90 min. The inoculum was then removed and replaced with 1.5 ml V medium, and cultures were incubated for a further 16 hours. Infected cells were harvested by removing the medium, washing the monolayers once with phosphate-buffered saline (PBS), scraping the cells into 1 ml PBS and pelleting the cells at 3,000 rpm in the microcentrifuge for 3 minutes. The supernatant liquid was removed, and cell lysates were prepared by resuspending the cell pellets in 25 µl water, adding 25 µl of 2X SDS-polyacrylamide gel sample buffer, and then incubating in a boiling water bath for 10 minutes. Proteins were separated by SDS-PAGE, blotted to nitrocellulose, and then probed with chicken antibody to UL34 (diluted 1:250) ^9^, mouse antibody to HSV-1 VP5 (diluted 1:500 (Biodesign International) mouse antibody to FLAG epitope (diluted 1:1000) (monoclonal M2, SIGMA/Aldrich) or rabbit antibody to calnexin (diluted 1:1000) (Cell Signaling Technology).

### Crystallization and data collection

Crystals of NEC185-Δ50-SUP_UL31_ were grown by vapor diffusion at 25 °C in hanging drops with 1 μL of protein (3 mg/mL), 1 μL of reservoir solution (10% PEG3350, 8 mM Li_2_SO_4_, 6 mM ATP, and 0.1 M MES, pH 6) and 1 μL of Silver Bullets (Hampton Research) reagent [G3 (0.25% 2,2’-Thiodiglycolic acid, 0.2% Azelaic acid, 0.2% Mellitic acid, 0.2% trans-aconitic acid, 0.02 M HEPES sodium pH 6.8)]. Hexagonal SUP_UL31_ crystals appeared after 2 days, only in the presence of Silver Bullets, and were completely grown after one week. Crystals were flash-frozen into liquid nitrogen in a solution identical to the reservoir solution containing 30% glycerol as the cryoprotectant.

Crystals of NEC185-Δ50-DN_UL34_/SUP_UL31_ were grown by vapor diffusion at 25 °C in hanging drops with 1 μL of protein (3.5 mg/mL), 1 μL of reservoir solution (10% PEG3350, 14 mM Li2SO4, 14 mM ATP, 10 mM phenol, and 0.1 M MES, pH 6) and 1 μL of Silver Bullets (Hampton Research) reagent [G3 (0.25% 2,2’-Thiodiglycolic acid, 0.2% Azelaic acid, 0.2% Mellitic acid, 0.2% trans-aconitic acid, 0.02 M HEPES sodium pH 6.8)]. Hexagonal DN_UL34_/SUP_UL31_ crystals appeared after one week, only in the presence of Silver Bullets. Crystals were flash-frozen into liquid nitrogen in a solution identical to the reservoir solution containing 30% glycerol as the cryoprotectant. In comparison to the SUP_UL31_ crystals, DN_UL34_/SUP_UL31_ crystals required an additional additive, phenol, and took longer to appear (one week vs. two days).

### Cryo-EM grid preparation and image collection

A volume of 10 μL of a 1:1 mixture of 400 nm and 800 nm large unilamellar vesicles (LUVs) made of 60% POPC/10% POPS/10% POPA/10% POPE/5% cholesterol/5% Ni-DGS [prepared as previously described ^17^] were mixed at room temperature with NEC220-SE6_UL31_-His_8_ to a final protein concentration of 1 mg/mL. After 15 min incubation, 3 μL of the sample was applied to glow-discharged (45 s) Quantifoil (2/2; Electron Microscopy Sciences) grids. Grids were blotted on both sides for 6 s with 0 blotting force and vitrified immediately by plunge freezing into liquid nitrogen-cooled liquid ethane (Vitrobot), before storage in liquid nitrogen. Grids were loaded into a Tecnai F20 transmission electron microscope (FEI) with a FEG and Compustage, equipped with a Gatan Oneview CMOS camera, using a cryo holder (Gatan) (Brandeis University Electron Microscope Facility). The microscope was operated in low-dose mode at 200 keV with SerialEM. Images were recorded at 19,000-fold (pixel size: 5.6 nm) magnification at a defocus of −4 μm. Images are displayed using ImageJ ^40^.

### Cryo-ET grid preparation

A volume of 10 μL of a 1:1 mixture of 400 nm and 800 nm large unilamellar vesicles (LUVs) made of POPC:POPS:POPA=3:1:1 [prepared as previously described ^17^] was mixed on ice with either WT-NEC220, NEC-SUP_UL31_, or NEC-DN_UL34_/SUP_UL31_, each to a final protein concentration of 1 mg/mL. After 30 min incubation, the sample was mixed with 5 nm fiducial gold beads, and cryo-ET grids were prepared by applying 3 μL of sample to glow-discharged (30 s) Lacey carbon grids (Electron Microscopy Sciences). Grids were blotted on both sides for 6 s with 0 blotting force and vitrified immediately by plunge freezing into liquid nitrogen-cooled liquid ethane with an FEI Mark IV Vitrobot cryo-sample plunger. Vitrified cryo-ET grids were stored in a liquid nitrogen dewar until use.

### Cryo-ET data collection and tomogram reconstruction

Tilt series were collected a Titan Krios electron microscope at the California NanoSystems Institute (CNSI). Data collection parameters are listed in **Supplementary Table 11**. Tilt series were collected using SerialEM ^41^ in a Titan Krios instrument equipped with a Gatan imaging filter (GIF) and a post-GIF K3 direct electron detector in electron-counting mode. Frames in each movie of the raw tilt series were aligned, drift-corrected, and averaged with Motioncor2 ^41^. The tilt series micrographs were aligned and reconstructed into 3D tomograms using the IMOD software package ^42^, then missing-wedge corrected by IsoNet ^43^ for particle picking.

### Sub-tomographic averaging

The variation in curvature of the NEC hexagonal coat made it difficult to identify the hexagonal repeat units required for particle picking. To overcome this, particle picking was performed using Python scripts derived from the Particle Estimation for Electron Tomography (PEET) software ^44^. Firstly, an initial model was generated as previously described ^45^ by manually picking ∼100 particles and performing sub-tomogram averaging using PEET. This allowed for the hexagonal geometric parameter, including the repeating distance and orients, of the NEC lattice to be accurately measured. Secondly, for each tomogram, a small set of particles were manually picked as “seed” particles sparsely covering all areas containing NEC. The “seed” particle set was then expanded by adding unknown particles near each of the known particles based on the hexagonal geometry obtained above. PEET alignment was performed on the expanded particles to match local conformational changes. Finally, the particle set expansion and PEET alignment were performed iteratively to obtain a complete particle set. Particles with less than three neighbors were excluded from the final particle set to remove outliers. Coordinates and orientations of the final particle set were formatted and imported into Relion ^46^ for further processing. One round of 3D refinement under bin4 pixel size and several rounds of 3D refinement and classification under bin2 pixel size, along with duplicate removal, resulted in the final masked resolutions: WT NEC (5.9 Å), NEC-DN_UL34_/SUP_UL31_ (5.4 Å), and NEC-SUP_UL31_ (13.1 Å). The resolutions reported above for the averaged structures are based on the ‘gold standard’ refinement procedures and the 0.143 Fourier shell correlation (FSC) criterion (**Fig. S5).**

### 3D Visualization

UCSF ChimeraX ^47^ was used to visualize the resulting sub-tomogram averages in their three dimensions, segmentation of density maps, and surface rendering for the different components of NEC.

## Data availability

All data generated or analyzed during this study are included in the manuscript and supporting files. A source data file is provided for data presented in Figs. 1-4. The cryo-ET sub-tomogram average maps have been deposited in the EM Data Bank under the accession codes EMD-40223 (WT NEC), EMD-40224 (NEC-DN_UL34_/SUP_UL31_), and EMD-40225 (NEC-SUP_UL31_). Atomic coordinates and structure factors for the NEC-SUP_UL31_ crystal structure have been deposited in the RCSB Protein Data Bank under accession code 8G6D. All files will be made publicly available upon publication.

## Acknowledgments

We thank Janna Bigalke (Tufts University) for cloning the SUP_UL31_, SE6_UL31_, and UL34 15-185 constructs; Xuanzong Guo (Tufts University) for cloning the UL31Δ50-306 SUP plasmid and for purifying the NEC-R139A_UL34_/SUP_UL31_ and NEC-D35A_UL34_/SUP_UL31_ constructs. We also thank the staff at the NE-CAT (Advanced Photon Source) for help with collecting x-ray diffraction data; Peter Cherepanov (Francis Crick Institute) for the gift of the GST-PreScission Protease expression plasmid; and Thomas Schwartz (Massachusetts Institute of Technology) for the gift of LoBSTr cells. We thank Rob Jackson (Tufts University School of Medicine) and Martin Hunter (University of Massachusetts College of Engineering) for help with fluorescence microscopy experiments. Confocal microscopy was performed at the Imaging and Cell Analysis Core Facility within the Center for Neuroscience Research at Tufts University School of Medicine, which is supported by NIH grant P30 NS047243 (Rob Jackson). Cryo-EM samples were prepared and imaged at the Brandeis Electron Microscopy Facility. This work is based upon research conducted at the Northeastern Collaborative Access Team beamlines, which are funded by the National Institute of General Medical Sciences from the National Institutes of Health (P30 GM124165). The Eiger 16M detector on the 24-ID-E beamline is funded by an NIH-ORIP HEI grant (S10OD021527). This research used resources of the Advanced Photon Source; a U.S. Department of Energy (DOE) Office of Science User Facility operated for the DOE Office of Science by Argonne National Laboratory under Contract No. DE-AC02-06CH11357. All software was installed and maintained by SBGrid ^48^. This work was supported by NIH grants R01GM111795 (E.E.H), R01AI147625 (E.E.H), R01DE028583 (Z.H.Z.), R01AI150718 (R.J.R), R21AI148831 (R.J.R), F32GM126760 (E.B.D), K12GM133314 (E.B.D), K99AI151891 (E.B.D), and by a Faculty Scholar grant 55108533 from Howard Hughes Medical Institute (E.E.H.).

## Author contributions

E.B.D and M.W. cloned, expressed, purified, and performed the corresponding in-vitro budding assays on the NEC constructs. R.J.R. performed the infected cell experiments including cloning and *trans*-complementation assays. E.B.D. crystallized NEC-SUP_UL31_ and NEC-DN_UL34_/SUP_UL31_. G.L.G. harvested crystals and assisted in data collection and processing. E.B.D. solved the NEC-SUP_UL31_ structure. H.W. collected, processed, and refined the NEC-WT, NEC-SUP_UL31_, and NEC-DN_UL34_/SUP_UL31_ cryo-ET data. S.L. assisted in cryo-ET data processing and averaging. R.J.R., Z.H.Z., and E.E.H. oversaw all aspects of the project. E.B.D. and E.E.H. wrote the manuscript. All authors edited and finalized the manuscript.

