## Supplementary Information for "The universal suppressor mutation in the HSV-1 nuclear egress complex restores membrane budding defects by stabilizing the oligomeric lattice"

Supplementary Figures S1-S5

Supplementary Tables S1-S18

Supplementary References

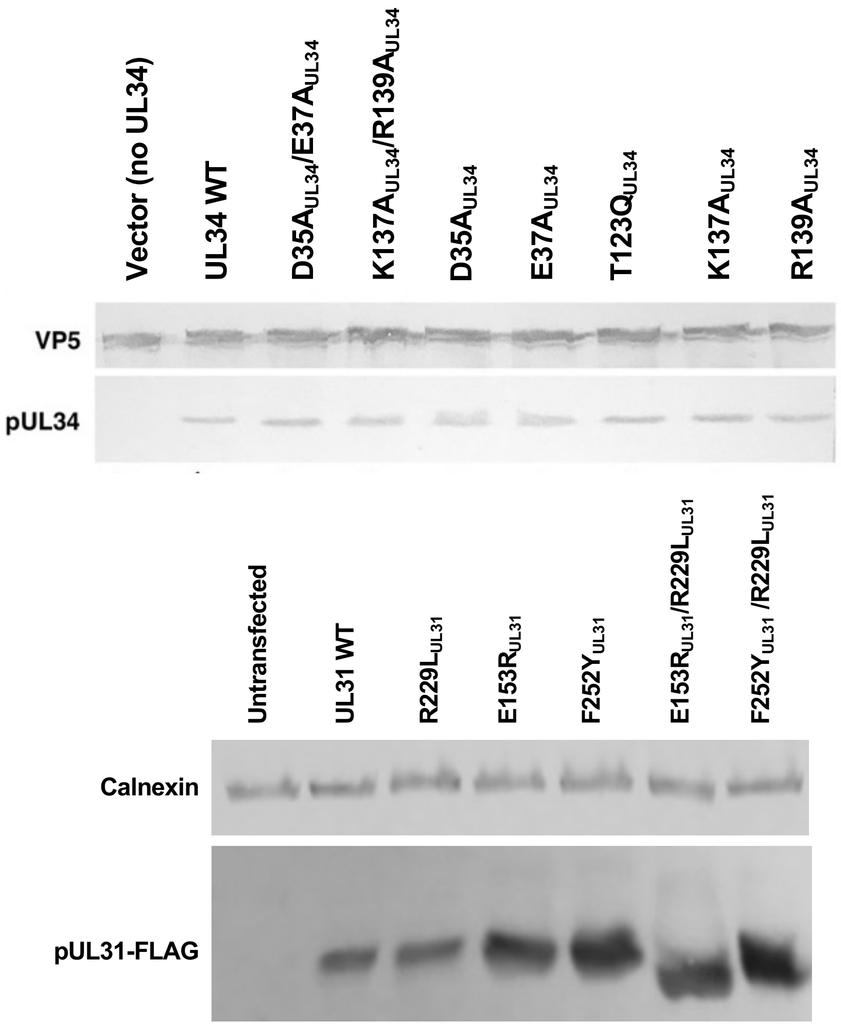

**Supplementary Figure 1.** Expression levels of WT UL31, WT UL34 and corresponding mutant proteins in Hep-2 cells used for *trans* complementation assays presented in this study.

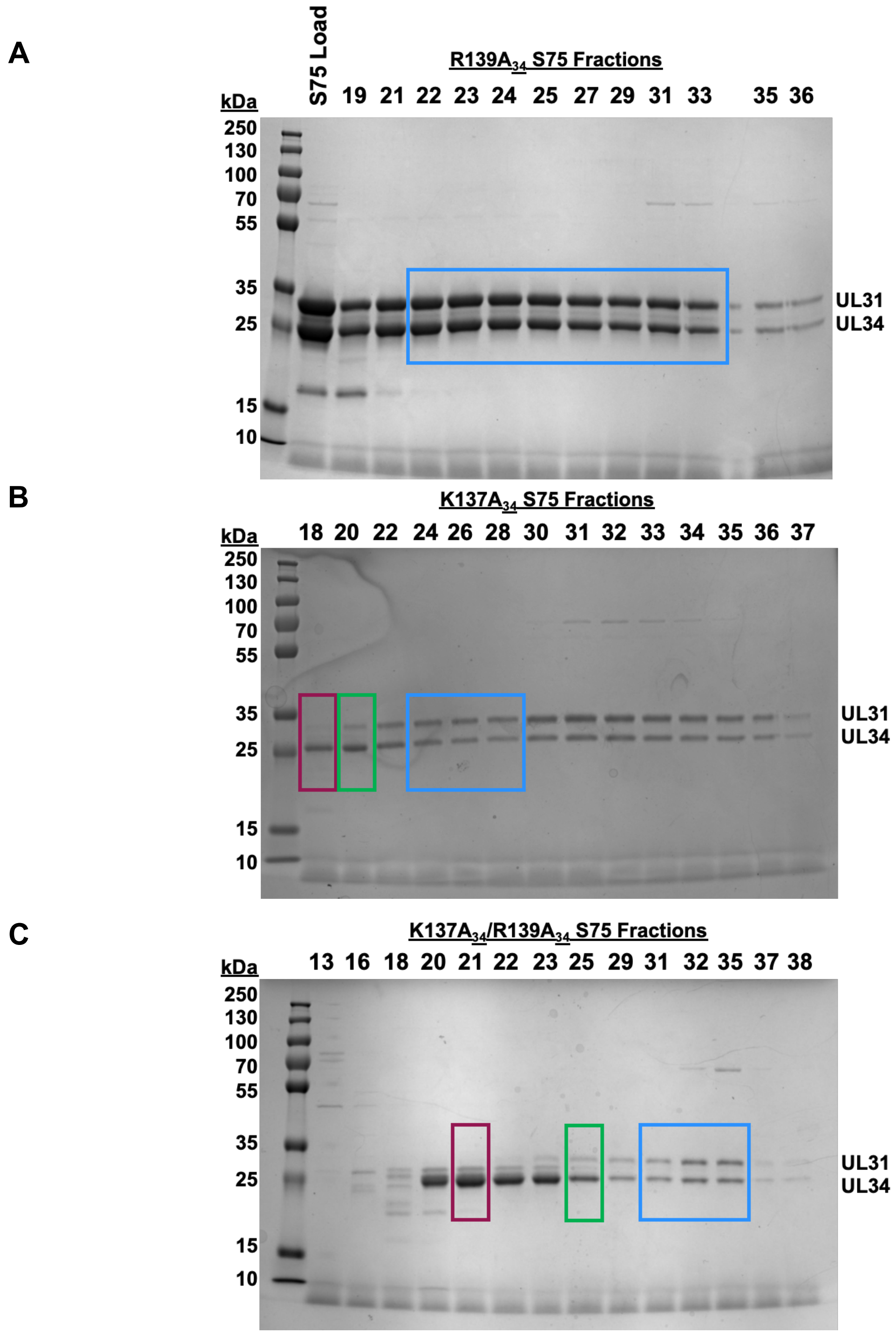

**Supplementary Figure 2.** SDS gels of **a)** NEC-R139A$_{\text{UL34}}$, **b)** NEC-K137A$_{\text{UL34}}$, and **c)** NEC-K137A$_{\text{UL34}}$/R139A$_{\text{UL34}}$. S75 purifications showing fractions containing either free UL34 (magenta), unequal amounts of UL31 and UL34 (green) and equimolar amounts of UL31 and UL34 (blue). Only fractions such as those in blue were collected for downstream studies. UL31: 34 kDa and UL34: 25 kDa.

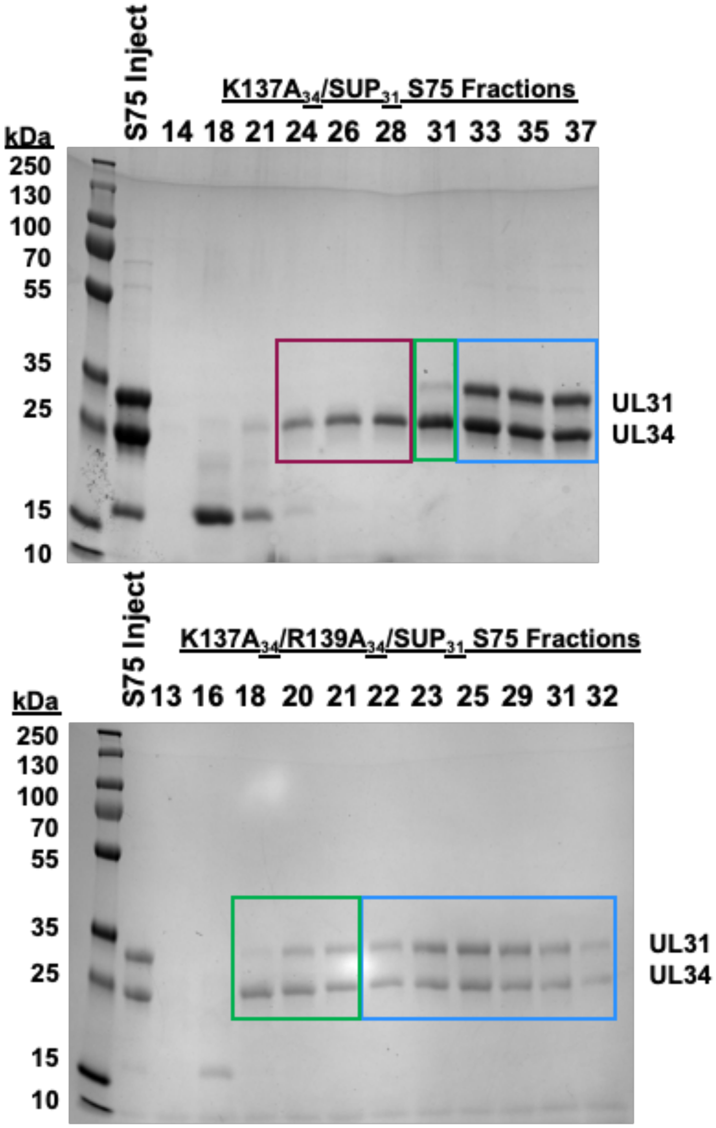

**Supplementary Figure 3.** SDS gels of NEC-K137A$_{UL34}$/SUP$_{UL31}$ and NEC-K137A$_{UL34}$/R139A$_{UL34}$/SUP$_{UL31}$ S75 purifications showing fractions containing either free UL34 (magenta), unequal amounts of UL31 and UL34 (green) or equimolar amounts of UL31 and UL34 (blue). Only fractions such as those in blue were collected for downstream studies. UL31: 34 kDa and UL34: 25 kDa.

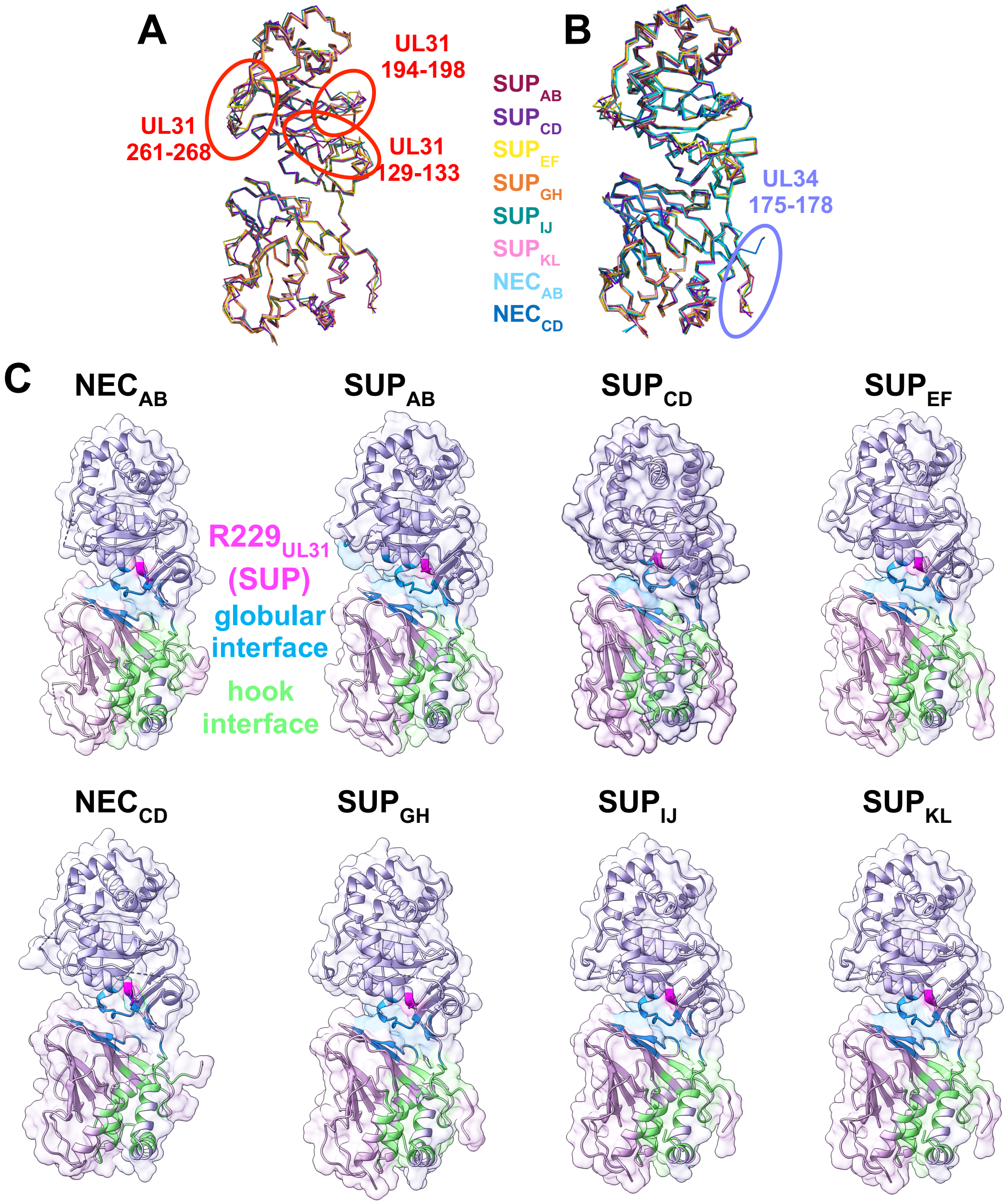

**Supplementary Figure 4. The six mutant SUP$_{UL31}$ heterodimers are similar to each other and to both the WT-NEC$_{AB}$ and WT-NEC$_{CD}$ heterodimers. a)** Backbone overlay of the six mutant SUP$_{UL31}$ heterodimers. Red circles indicate variable regions in UL31: residues 194-198, 128-136, and 260-266. **b)** Backbone overlay of WT-NEC$_{AB}$ and WT-NEC$_{CD}$ to the six SUP$_{UL31}$ heterodimers. The purple circle indicates additional UL34 C-terminus residues (175-178) resolved in the SUP$_{UL31}$ heterodimers that were unresolved in the WT structures. **c)** Comparison of the UL31/UL34 heterodimeric interfaces: globular core (blue) and UL31 hook (green). The position of either R229 in the WT structures or L229 in the SUP structures is also shown (magenta).

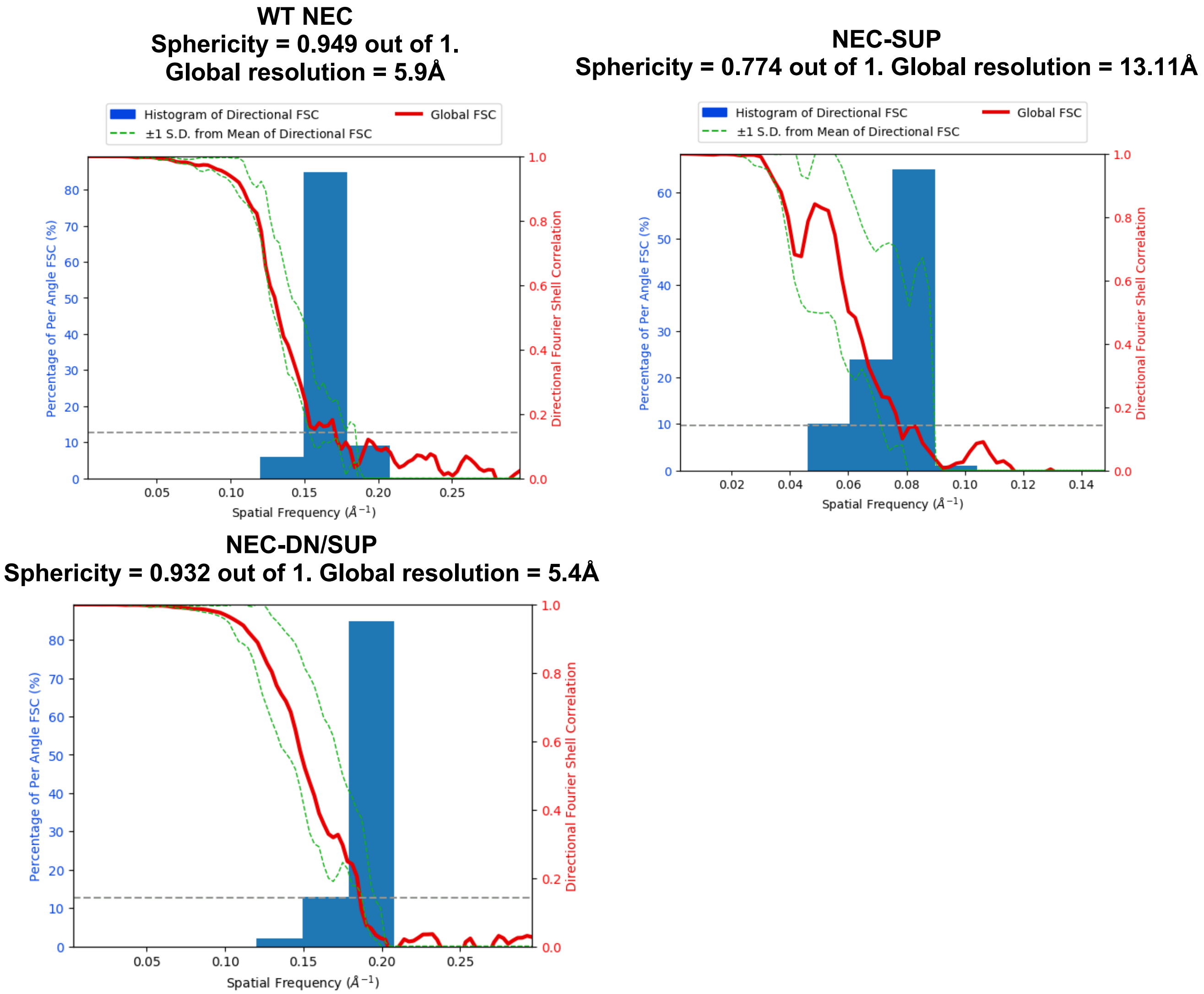

**Supplementary Figure 5.** Directional Fourier shell correlation (FSC) curves [1] for the subtomogram averages of NEC lattice for WT NEC, DNSUP-NEC, and SUP-NEC.

**Supplementary Table S1. Data collection and refinement statistics for NEC-SUPUL31.**

| Parameter | Value[a] |
|---|---|
| *Data collection statistics* | |
| Wavelength (Å) | 0.9786 |
| Space group | $C12_1$ |
| Unit cell | $a$ = 109.555 Å, $b$ = 189.103 Å, $c$ = 157.096 Å, $\alpha = \gamma = 90°$, $\beta$ = 100.52° |
| Resolution range (Å) | 94.55 – 3.92 (4.06 – 3.92) |
| No. of reflections | |
| *Total* | 189091 (18304) |
| *Unique* | 28047 (2801) |
| Multiplicity | 6.7 (6.5) |
| Completeness (%) | 99.07 (98.83) |
| Mean $I/\sigma(I)$ | 2.83 (1.51) |
| Wilson B-factor (Å$^2$) | 59.92 |
| $R_{merge}$ | 1.84 (6.109) |
| $R_{meas}$ | 1.996 (6.637) |
| $R_{pim}$ | 0.7632 (2.564) |
| $CC_{1/2}$ | 0.591 (0.312) |
| CC* | 0.862 (0.689) |
| *Refinement Statistics* | |
| No. of reflections used | |
| *In refinement* | 27993 (2795) |
| *For $R_{free}$* | 1996 (199) |
| $R_{work}$[b] | 0.2546 (0.3051) |
| $R_{free}$[b] | 0.3011 (0.3443) |
| $CC_{work}$ | 0.873 (0.776) |
| $CC_{free}$ | 0.865 (0.750) |
| *No of:* | |
| Nonhydrogen atoms | 19341 |
| Macromolecules | 19333 |
| Ligands | 6 |
| Solvent | 2 |
| Protein Residues | 2492 |
| *RMSD* | |
| Bond length (Å) | 0.0003 |
| Bond angle (°) | 0.70 |
| *Ramachandran plot[c] (%)* | |
| Favored regions | 93.67 |
| Allowed regions | 5.72 |
| Outliers | 0.61 |
| Rotamer outliers (%) | 0.00 |
| Clash score | 8.33 |
| *B-factor* | |

| | |
|---|---|
| Average | 70.14 |
| Macromolecules | 70.13 |
| Ligands | 119.26 |
| Solvent | 39.49 |

[a] Highest resolution shell statistics are shown in parentheses.
[b] $R_{work}$ and $R_{free}$ are defined as $\sum||F_{obs}| - |F_{calc}||/\sum|F_{obs}|$ for the reflections in the working or the test set, respectively.
[c] Determined using MolProbity[2].

**Supplementary Table S2. Resolved residues for each chain of UL34 (top) and UL31 (bottom) from the NEC-SUP$_{UL31}$ crystal structure.** Resolved residue boundaries, along with % resolved, are provided for each of the chains.

| **Chain ID** | **Resolved Residues** | **% Resolved Residues** |
|---|---|---|
| **UL34** | | **Total Residues: 171** |
| **A** | 15-178 | 96 |
| **C** | 15-178 | 96 |
| **E** | 15-178 | 96 |
| **G** | 15-176 | 94 |
| **I** | 15-176 | 94 |
| **K** | 15-177 | 95 |
| **UL31** | | **Total Residues: 255** |
| **B** | 54-306 | 99 |
| **D** | 54-306 | 99 |
| **F** | 54-306 | 99 |
| **H** | 54-131, 135-306 | 96 |
| **J** | 57-306 | 99 |
| **L** | 54-128, 134-306 | 96 |

**Supplementary Table S3. Structural alignments of the NEC-SUP$_{UL31}$ heterodimers in the asymmetric unit.** NEC-SUP$_{UL31}$ heterodimers and individual UL34 and UL31 chains were aligned. RMSD (Å) values are reported and were calculated using "SSM Superpose" in WinCoot [3].

| | **UL34$_A$/ UL31$_B$** | **UL34$_C$/ UL31$_D$** | **UL34$_E$/ UL31$_F$** | **UL34$_G$/ UL31$_H$** | **UL34$_I$/ UL31$_J$** | **UL34$_K$/ UL31$_L$** |
|---|---|---|---|---|---|---|
| **UL34$_A$/ UL31$_B$** | -- | 0.91 | 0.85 | 0.86 | 0.99 | 0.69 |
| **UL34$_C$/ UL31$_D$** | 0.91 | -- | 0.85 | 0.85 | 0.96 | 0.85 |
| **UL34$_E$/ UL31$_F$** | 0.85 | 0.85 | -- | 0.73 | 0.94 | 0.72 |
| **UL34$_G$/ UL31$_H$** | 0.86 | 0.85 | 0.73 | -- | 0.97 | 0.67 |
| **UL34$_I$/ UL31$_J$** | 0.99 | 0.96 | 0.94 | 0.97 | -- | 1.00 |
| **UL34$_K$/ UL31$_L$** | 0.69 | 0.85 | 0.72 | 0.67 | 1.00 | -- |
| **UL34** | **Chain A** | **Chain C** | **Chain E** | **Chain G** | **Chain I** | **Chain K** |
| **Chain A** | -- | 0.72 | 0.64 | 0.75 | 0.69 | 0.68 |
| **Chain C** | 0.72 | -- | 0.73 | 0.55 | 0.75 | 0.73 |
| **Chain E** | 0.64 | 0.73 | -- | 0.56 | 0.63 | 0.59 |
| **Chain G** | 0.75 | 0.55 | 0.56 | -- | 0.58 | 0.54 |
| **Chain I** | 0.69 | 0.75 | 0.63 | 0.58 | -- | 0.63 |
| **Chain K** | 0.68 | 0.73 | 0.59 | 0.54 | 0.63 | -- |
| **UL31** | **Chain B** | **Chain D** | **Chain F** | **Chain H** | **Chain J** | **Chain L** |
| **Chain B** | -- | 0.90 | 0.82 | 0.84 | 1.04 | 0.64 |
| **Chain D** | 0.90 | -- | 0.89 | 0.93 | 1.02 | 0.85 |
| **Chain F** | 0.82 | 0.93 | -- | 0.80 | 1.00 | 0.69 |
| **Chain H** | 0.84 | 0.93 | 0.80 | -- | 1.01 | 0.70 |
| **Chain J** | 1.04 | 1.02 | 1.00 | 1.01 | -- | 1.03 |
| **Chain L** | 0.64 | 0.85 | 0.69 | 0.70 | 1.03 | -- |

**Supplementary Table 4. Structural alignments of the NEC-SUP$_{UL31}$ and WT-NEC heterodimers.** NEC- SUP$_{UL31}$ heterodimers and individual UL34 and UL31 chains were aligned with the two WT NEC heterodimers, along with the corresponding WT individual chains. RMSD (Å) values are reported and were calculated using "SSM Superpose" in WinCoot [3].

| **SUP** | **WT UL34$_A$/UL31$_B$** | **WT UL34$_C$/UL31$_D$** |
|---|---|---|
| **UL34$_A$/UL31$_B$** | 0.94 | 0.91 |
| **UL34$_C$/UL31$_D$** | 0.99 | 1.02 |
| **UL34$_E$/UL31$_F$** | 0.84 | 0.82 |
| **UL34$_G$/UL31$_H$** | 0.93 | 0.90 |
| **UL34$_I$/UL31$_J$** | 0.85 | 0.83 |
| **UL34$_K$/UL31$_L$** | 0.91 | 0.95 |
| **SUP** | **WT UL34$_A$** | **WT UL34$_C$** |
| **UL34$_A$** | 0.72 | 0.68 |
| **UL34$_C$** | 0.82 | 0.87 |
| **UL34$_E$** | 0.71 | 0.74 |
| **UL34$_G$** | 0.72 | 0.76 |
| **UL34$_I$** | 0.68 | 0.75 |
| **UL34$_K$** | 0.68 | 0.80 |
| **SUP** | **WT UL31$_B$** | **WT UL31$_D$** |
| **UL31$_B$** | 0.91 | 0.86 |
| **UL31$_D$** | 0.97 | 0.98 |
| **UL31$_F$** | 0.82 | 0.73 |
| **UL31$_H$** | 0.81 | 0.87 |
| **UL31$_J$** | 0.82 | 0.84 |
| **UL31$_L$** | 0.87 | 0.91 |

**Supplementary Table S5. Residues involved in heterodimeric interactions in the WT-NEC UL34$_A$/UL31$_B$, WT-NEC UL34$_C$/UL31$_D$, and the SUP$_{UL31}$ mutant heterodimers.** Interface residues between UL31 and UL34 hook interface 1 (boxes shaded light green) and globular interface 2 (boxes shaded in blue) were determined using PDBePISA analysis [4]. Residues that were not resolved in the structure are indicated with NR.

| | Residue | WT UL34$_A$/UL31$_B$ | WT UL34$_C$/UL31$_D$ | SUP UL34$_A$/UL31$_B$ | SUP UL34$_C$/UL31$_D$ | SUP UL34$_E$/UL31$_F$ | SUP UL34$_G$/UL31$_H$ | SUP UL34$_I$/UL31$_J$ | SUP UL34$_K$/UL31$_L$ |
|---|---|---|---|---|---|---|---|---|---|
| UL31 Residues | C54 | NR | NR | | | | | NR | |
| | L55 | | | | | | | NR | |
| | H56 | | | | | | | NR | |
| | E57 | | | | | | | | |
| | R58 | | | | | | | | |
| | Q59 | | | | | | | | |
| | R60 | | | | | | | | |
| | Y61 | | | | | | | | |
| | R62 | | | | | | | | |
| | L64 | | | | | | | | |
| | F65 | | | | | | | | |
| | L68 | | | | | | | | |
| | A69 | | | | | | | | |
| | P72 | | | | | | | | |
| | D74 | | | | | | | | |
| | E75 | | | | | | | | |
| | I76 | | | | | | | | |
| | I78 | | | | | | | | |
| | V79 | | | | | | | | |
| | R80 | | | | | | | | |
| | S81 | | | | | | | | |
| | L82 | | | | | | | | |
| | S83 | | | | | | | | |
| | V84 | | | | | | | | |
| | P85 | | | | | | | | |
| | L86 | | | | | | | | |
| | V87 | | | | | | | | |
| | T90 | | | | | | | | |
| | P91 | | | | | | | | |
| | D99 | | | | | | | | |
| | Q100 | | | | | | | | |
| | T101 | | | | | | | | |
| | V102 | | | | | | | | |
| | A103 | | | | | | | | |
| | D104 | | | | | | | | |
| | N105 | | | | | | | | |

| | |
|---|---|
| | **L116** |
| | **I118** |
| | **D232** |
| | **T236** |
| | **K262** |
| **UL34 Residues** | **R22** |
| | **L25** |
| | **I26** |
| | **V27** |
| | **P28** |
| | **P65** |
| | **D67** |
| | **Y68** |
| | **R71** |
| | **L72** |
| | **N74** |
| | **D75** |
| | **A77** |
| | **E78** |
| | **P80** |
| | **C81** |
| | **N82** |
| | **P83** |
| | **E114** |
| | **R115** |
| | **T116** |
| | **N117** |
| | **V118** |
| | **I119** |
| | **L130** |
| | **G131** |
| | **D134** |
| | **K137** |
| | **L140** |
| | **L142** |
| | **A144** |
| | **P146** |
| | **M147** |
| | **A149** |
| | **S150** |
| | **W152** |
| | **F156** |
| | **R158** |

| | |
|---|---|
| | R161 |
| | Q163 |
| | L164 |
| | A165 |
| | R167 |
| | F168 |
| | M169 |
| | G170 |
| | P171 |
| | D172 |
| | G175 |

**Supplementary Table S6. Residues involved in hexameric interactions in the WT-NEC $UL34_A/UL31_B$ lattice, WT-NEC $UL34_C/UL31_D$ lattice, and the SUP heterodimers within the SUP lattice.** Interface residues between UL31 and UL34 (boxes shaded in teal) and between UL34 and UL34 (boxes shaded in dark green) were determined using PDBePISA analysis [4]. Residues that were not resolved in the structure are indicated with NR.

| | | WT $UL34_A/UL31_B$ or $UL34_A/UL34_A$ | WT $UL34_C/UL31_D$ or $UL34_C/UL34_C$ | SUP $UL34_A/UL31_J$ or $UL34_A/UL34_I$ | SUP $UL34_G/UL31_L$ or $UL34_G/UL34_K$ | SUP $UL34_I/UL31_F$ or $UL34_I/UL34_E$ | SUP $UL34_E/UL31_D$ or $UL34_E/UL34_C$ | SUP $UL34_C/UL31_H$ or $UL34_C/UL34_G$ | SUP $UL34_K/UL31_B$ or $UL34_K/UL34_A$ |
|---|---|---|---|---|---|---|---|---|---|
| UL31 Residues | V87 | | | | | | | | |
| | T89 | | | | | | | | |
| | P91 | | | | | | | | |
| | L94 | | | | | | | | |
| | S110 | | | | | | | | |
| | G111 | | | | | | | | |
| | M112 | | | | | | | | |
| | G113 | | | | | | | | |
| | Y114 | | | | | | | | |
| | Y115 | | | | | | | | |
| | T222 | | | | | | | | |
| | H246 | | | | | | | | |
| | V247 | | | | | | | | |
| | Q249 | | | | | | | | |
| | S250 | | | | | | | | |
| | F252 | | | | | | | | |
| UL34 Residues | G33 | | | | | | | | |
| | G34 | | | | | | | | |
| | D35 | | | | | | | | |
| | E37 | | | | | | | | |
| | Y41 | | | | | | | | |
| | S45 | | | | | | | | |

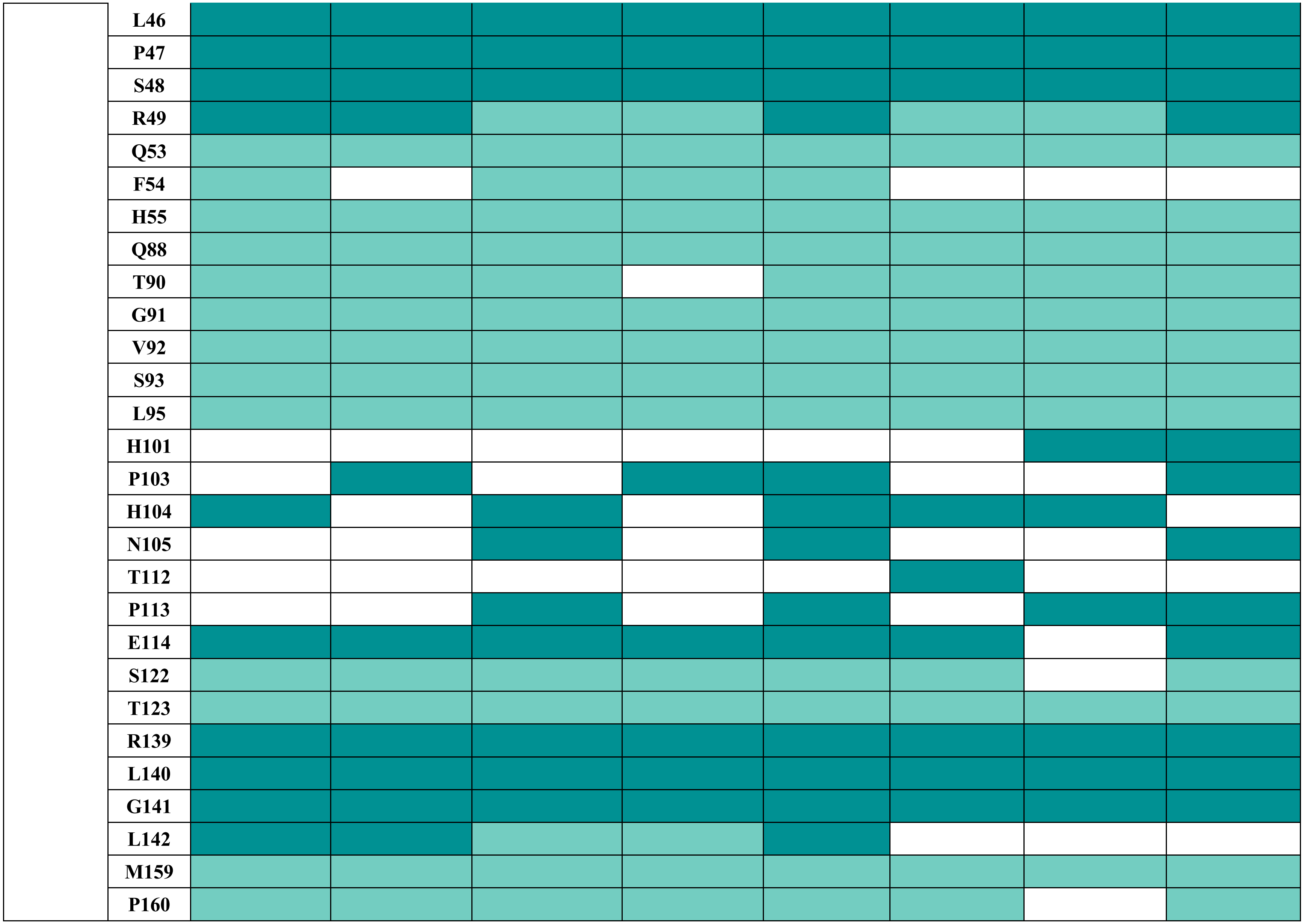
L46
P47
S48
R49
Q53
F54
H55
Q88
T90
G91
V92
S93
L95
H101
P103
H104
N105
T112
P113
E114
S122
T123
R139
L140
G141
L142
M159
P160

**Supplementary Table S7. Conservation of residues at the hexameric interfaces within the NEC-SUP crystal structure compared to the two types of WT-NEC lattices (A/B and C/D).** Interface residues were determined using PDBePISA analysis [4].

| | **WT A/B: $UL34_A/UL31_B$ and $UL34_A/UL34_A$ (42 resolved hexameric interface residues)** | | |
|---|---|---|---|
| **SUP Hexameric Interface** | Identical Interface Residues | Total interface residues | % Identical interface residues (compared to WT) |
| **$UL34_A/UL31_J$ and $UL34_A/UL34_I$** | 35 | 40 | 88 |
| **$UL34_G/UL31_L$ and $UL34_G/UL34_K$** | 34 | 37 | 92 |
| **$UL34_I/UL31_F$ and $UL34_I/UL34_E$** | 35 | 38 | 92 |
| **$UL34_E/UL31_D$ and $UL34_E/UL34_C$** | 33 | 35 | 94 |
| **$UL34_C/UL31_H$ and $UL34_C/UL34_G$** | 29 | 32 | 91 |
| **$UL34_K/UL31_B$ and $UL34_K/UL34_A$** | 40 | 42 | 95 |
| | **WT C/D: $UL34_C/UL31_D$ and $UL34_C/UL34_C$ (39 resolved hexameric interface residues)** | | |
| **SUP Hexameric Interface** | Identical Interface Residues | Total interface residues | % Identical interface residues (compared to WT) |
| **$UL34_A/UL31_J$ and $UL34_A/UL34_I$** | 33 | 40 | 85 |
| **$UL34_G/UL31_L$ and $UL34_G/UL34_K$** | 36 | 37 | 97 |
| **$UL34_I/UL31_F$ and $UL34_I/UL34_E$** | 35 | 38 | 92 |
| **$UL34_E/UL31_D$ and $UL34_E/UL34_C$** | 33 | 35 | 94 |
| **$UL34_C/UL31_H$ and $UL34_C/UL34_G$** | 28 | 32 | 88 |
| **$UL34_K/UL31_B$ and $UL34_K/UL34_A$** | 37 | 42 | 95 |

**Supplementary Table S8. Conservation of residues at the interhexameric interfaces within the NEC-SUP crystal structure compared to the two types of WT-NEC lattices (A/B and C/D).** Interface residues were determined using PDBePISA analysis [4].

| **Interhexameric Interface** | | **WT UL31$_{B}$/UL31$_{B}$/UL31$_{B}$ (13 total resolved)** | | |
|---|---|---|---|---|
| | | Identical interface residues | Total resolved SUP interface residues | % Identical interface residues (compared to WT) |
| **Trimer 1** (UL31 only) | **SUP UL31$_{B}$/UL31$_{H}$/UL31$_{F}$** | 13 | 26 | 100 |
| **Trimer 2** (UL31 only) | **SUP UL31$_{D}$/UL31$_{J}$/UL31$_{L}$** | 12 | 20 | 92 |
| | | **WT UL31$_{D}$/UL31$_{D}$/UL31$_{D}$ (7 total resolved)** | | |
| **Trimer 1** (UL31 only) | **SUP UL31$_{B}$/UL31$_{H}$/UL31$_{F}$** | 4 | 26 | 57 |
| **Trimer 2** (UL31 only) | **SUP UL31$_{D}$/UL31$_{J}$/UL31$_{L}$** | 3 | 20 | 43 |
| | | **WT UL34$_{A}$/UL31$_{B}$ and UL34$_{A}$/UL31$_{B}$ (7 total resolved)** | | |
| **Dimer 1** (UL31 and UL34) | **SUP UL34$_{A}$/UL31$_{B}$ and UL34$_{C}$/UL31$_{D}$** | 6 | 12 | 86 |
| | **SUP UL34$_{E}$/UL31$_{F}$ and UL34$_{K}$/UL31$_{L}$** | 6 | 7 | 86 |
| | **SUP UL34$_{G}$/UL31$_{H}$ and UL34$_{I}$/UL31$_{J}$** | 6 | 11 | 86 |
| **Dimer 2** (UL31 only) | **SUP UL31$_{B}$/UL31$_{D}$** | 4 | 4 | 100 |
| | **SUP UL31$_{F}$/UL31$_{L}$** | 4 | 4 | 100 |
| | **SUP UL31$_{H}$/UL31$_{J}$** | 3 | 3 | 100 |
| | | **WT UL34$_{C}$/UL31$_{D}$ and UL34$_{C}$/UL31$_{D}$ (20 total resolved)** | | |
| **Dimer 1** (UL31 and UL34) | **SUP UL34$_{A}$/UL31$_{B}$ and UL34$_{C}$/UL31$_{D}$** | 4 | 12 | 20 |
| | **SUP UL34$_{E}$/UL31$_{F}$ and UL34$_{K}$/UL31$_{L}$** | 4 | 7 | 20 |
| | **SUP UL34$_{G}$/UL31$_{H}$ and UL34$_{I}$/UL31$_{J}$** | 5 | 11 | 25 |
| **Dimer 2** (UL31 only) | **SUP UL31$_{B}$/UL31$_{D}$** | 0 | 4 | 0 |
| | **SUP UL31$_{F}$/UL31$_{L}$** | 0 | 4 | 0 |
| | **SUP UL31$_{H}$/UL31$_{J}$** | 0 | 3 | 0 |

**Supplementary Table S9. Residues involved in interhexameric (trimeric) interactions in the WT-NEC UL34$_A$/UL31$_B$ lattice, WT-NEC UL34$_C$/UL31$_D$ lattice, and the SUP heterodimers within the SUP lattice.** Interface residues (boxes shaded in light orange) were determined using PDBePISA analysis [4]. Unresolved residues are indicated with NR.

| | | WT UL31$_B$/UL31$_B$/ UL31$_B$ | WT UL31$_D$/UL31$_D$/ UL31$_D$ | SUP UL31$_B$/UL31$_H$/UL31$_F$ (Trimer 1) | | | SUP UL31$_D$/UL31$_J$/UL31$_L$ (Trimer 2) | | |
|---|---|---|---|---|---|---|---|---|---|
| | | | | **B** | **H** | **F** | **D** | **J** | **L** |
| UL31 Residues | **D129** | | NR | near | | | near | near | NR |
| | **G130** | | NR | | | | | | NR |
| | **R131** | NR | NR | | | | | | NR |
| | **F132** | NR | NR | | NR | | | | NR |
| | **A133** | | NR | | NR | | | | NR |
| | **A134** | | | | NR | | | | |
| | **S136** | | | | | | | | |
| | **E138** | | | | | | | | |
| | **A139** | | | | | | | | |
| | **I141** | | | | | | | | |
| | **L142** | | | | | | | | |
| | **V145** | | | | | | | | |
| | **Q146** | | | | | | | | |
| | **N149** | | | | | | | | |
| | **T150** | | | | | | | | |
| | **F152** | | | | | | | | |
| | **E153** | | | | | | | | |
| | **R155** | | | | | | | | |
| | **R193** | | | | | | | | |
| | **G194** | | | | | | | | |
| | **G195** | | | | | | | | |
| | **G196** | | | | | | | | |

| | | | |
|---|---|---|---|
| | **A197** | | |
| | **D199** | | |
| | **E267** | **NR** | **NR** |
| | **P269** | | |
| | **D286** | | |
| | **G287** | | |
| | **G288** | | |

**Supplementary Table S10. Residues involved in interhexameric (dimeric) interactions in the WT-NEC $UL34_A/UL31_B$ lattice, WT-NEC $UL34_C/UL31_D$ lattice, and the SUP heterodimers within the SUP lattice.** Interface residues are shaded in light orange (Dimer 1; UL31/UL31 and UL34/UL34) and dark orange (Dimer 2; UL31/UL31). Interfaces were determined using PDBePISA analysis [4].

| | | **WT $UL31_B/UL31_B$** | **WT $UL31_D/UL31_D$** | **SUP $UL31_B/UL31_D$** | | **SUP $UL31_F/UL31_L$** | | **SUP $UL31_H/UL31_J$** | |
|---|---|---|---|---|---|---|---|---|---|
| | | **B** | **D** | **B** | **D** | **F** | **L** | **H** | **J** |
| **UL31 Residues** | **P72** | | | | | | | | |
| | **S73** | | | | | | | | |
| | **E74** | | | | | | | | |
| | **I76** | | | | | | | | |
| | **A77** | | | | | | | | |
| | **S81** | | | | | | | | |
| | **N126** | | | | | | | | |
| | **S136** | | | | | | | | |
| | **E138** | | | | | | | | |
| | **A139** | | | | | | | | |
| | **I141** | | | | | | | | |
| | **L142** | | | | | | | | |
| | **V145** | | | | | | | | |
| | **Q146** | | | | | | | | |
| | **P269** | | | | | | | | |
| | **G287** | | | | | | | | |
| | **G288** | | | | | | | | |
| | **L291** | | | | | | | | |

| | | | | | | | | | |
|---|---|---|---|---|---|---|---|---|---|
| | **R295** | | | | | | | | |
| | | **WT**<br>**$UL34_A$/$UL34_A$** | **WT**<br>**$UL34_C$/$UL34_C$** | **SUP**<br>**$UL34_A$/$UL34_C$** | | **SUP**<br>**$UL34_E$/$UL34_K$** | | **SUP**<br>**$UL34_G$/$UL34_I$** | |
| | | | | **A** | **C** | **E** | **K** | **G** | **I** |
| **UL34 Residues** | **P14** | | | | | | | | |
| | **A15** | | | | | | | | |
| | **F16** | | | | | | | | |
| | **E17** | | | | | | | | |
| | **Q21** | | | | | | | | |
| | **R24** | | | | | | | | |
| | **L25** | | | | | | | | |
| | **R32** | | | | | | | | |
| | **G33** | | | | | | | | |
| | **D35** | | | | | | | | |
| | **H55** | | | | | | | | |
| | **H57** | | | | | | | | |
| | **E172** | | | | | | | | |
| | **D173** | | | | | | | | |
| | **A174** | | | | | | | | |

**Supplementary Table S11. Conditions used for cryo-ET data collection and processing of NEC and mutants presented in this study.**

| Cryo-ET Data Collection Statistics | | | |
|---|---|---|---|
| | **NEC-WT** | **NEC-$DN_{UL34}$/$SUP_{UL31}$** | **NEC-$SUP_{UL31}$** |
| **Data collection and processing** | | | |
| **Microscope** | Titan Krios | | |
| **Voltage (kV)** | 300 | | |
| **Total Electron exposure (e–/Å2)** | 110 | | |
| **Slit width (eV)** | 20 | | |
| **Detector** | K3 | | |
| **Defocus range (µm)** | -2.5 to -5.5 | | |
| **Pixel size (Å)** | 1.69 | | |
| **Software** | SerialEM [5] | | |
| **Tilt-series range** | ±46° | | |
| **Tilt-series increment** | ±2° | | |
| **Tilt-series scheme** | Dose symmetry | | |
| **EM grid type** | Lacey carbon | | |
| **Tilt-series used** | 43 | 35 | 2 |
| **Data processing** | | | |
| **Software: tilt-series alignment** | IMOD [6] | | |
| **Software: final reconstruction** | Relion 4.0 [7] | | |
| **Initial particle images (no.)** | 83385 | 48481 | 1564 |
| **Final particle images (no.)** | 34223 | 35039 | 1390 |
| **Final Box-size (pixel)** | $196^3$ | $196^3$ | $128^3$ |
| **Pixel size final reconstruction (Å)** | 1.69 | 1.69 | 3.38 |
| **Symmetry imposed** | C6 | | |
| **Masked map resolution (Å)** | 5.9 | 5.4 | 13.1 |
| **FSC threshold** | 0.143 | | |

**Supplementary Table S12. Buried surface areas at the hexameric interfaces.** PDBePISA analysis [4] was used to calculate the buried surface areas at the hexameric interfaces (either between UL31/UL34 or UL34/UL34) within the NEC-hexamer$_{AB}$, NEC-hexamer$_{CD}$, and SUP$_{UL31}$ mutant lattices. The total hexameric interface buried surface area was calculated by adding the UL31/UL34 and UL34/UL34 surface areas together. For WT HSV-1, the 4ZXS PDB structure was used.

| Protein | Chains at hexameric interface | UL31/UL34 interface area (Å) | UL34/UL34 interface area (Å) | Total hexameric interface area (Å) |
|---|---|---|---|---|
| **WT** | **UL34$_A$/UL31$_B$ and UL34$_A$/UL34$_A$** | 613 | 228 | 842 |
| | **UL34$_C$/UL31$_D$ and UL34$_C$/UL34$_C$** | 572 | 258 | 830 |
| **SUP** | **UL34$_A$/UL31$_J$ and UL34$_A$/UL34$_I$** | 614 | 179 | 793 |
| | **UL34$_G$/UL31$_L$ and UL34$_G$/UL34$_K$** | 619 | 186 | 805 |
| | **UL34$_I$/UL31$_F$ and UL34$_I$/UL34$_E$** | 531 | 225 | 756 |
| | **UL34$_E$/UL31$_D$ and UL34$_E$/UL34$_C$** | 550 | 185 | 735 |
| | **UL34$_C$/UL31$_H$ and UL34$_C$/UL34$_G$** | 526 | 189 | 715 |
| | **UL34$_K$/UL31$_B$ and UL34$_K$/UL34$_A$** | 560 | 282 | 842 |

**Supplementary Table S13. Comparison of contacts made at the WT NEC and NEC-SUP$_{UL31}$ lattice hexameric interfaces.** Atomic contacts (hydrogen bonds or salt bridges) between heterodimers within a hexameric lattice (shaded in blue) determined using PDBePISA [4].

| | UL31 Residue | UL34 Residue | WT A/B | WT C/D | SUP A/J | SUP L/G | SUP F/I | SUP D/E | SUP H/C | SUP B/K |
|---|---|---|---|---|---|---|---|---|---|---|
| H-bonds | Tyr 114 OH | Thr 90 O | ■ | ■ | ■ | | ■ | | ■ | ■ |
| | Tyr 114 OH | Thr 123 OG1 | | | ■ | | | | | |
| | Leu 86 O | Arg 49 NH1 | ■ | | | ■ | ■ | | | |
| | Leu 86 O | Arg 49 NH2 | ■ | ■ | | | | ■ | ■ | |
| | Lys 88 N | Glu 37 OE2 | | | | | | | | ■ |
| | Thr 89 OG1 | Glu 37 OE2 | | ■ | | ■ | | ■ | | |
| | Thr 89 OG1 | Glu 37 OE1 | | | | | ■ | | ■ | |
| | Thr 89 OG1 | Arg 49 NH2 | | | | ■ | | | | |
| | Thr 89 O | Arg 49 NH1 | | | ■ | | | | | |
| | Thr 89 N | Glu 37 OE2 | | | | | | | | ■ |
| | Gly 111 O | Arg 49 NH2 | ■ | | | | | | | |
| | Gly 111 O | Thr 90 OG1 | | | ■ | | | ■ | ■ | |
| | Gly 111 O | Ser 93 OG | | ■ | | | ■ | ■ | | |
| | Met 112 O | Gln 53 NE2 | ■ | ■ | | ■ | | | ■ | |
| | Met 112 SD | His 55 ND1 | | | ■ | | | | | |
| | Tyr 114 OH | Gly 91 N | | | ■ | ■ | | ■ | | |
| | Ser 250 OG | Gly 91 O | | | | ■ | | | | ■ |
| | Ser 250 OG | Met 159 O | | | | | | | ■ | |
| | **UL34 Residue** | **UL34 Residue** | **WT A/A** | **WT C/C** | **SUP A/K** | **SUP E/I** | **SUP G/C** | **SUP G/K** | **SUP C/E** | **SUP A/I** |
| H- bond | Glu 114 OE2 | His 104 N | ■ | | | | | | | |
| | Arg 139 O | Ser 48 N | ■ | ■ | | | | | | |
| | Leu 140 O | Ser 48 N | | ■ | | | | | | ■ |
| | Leu 140 O | Ser 48 OG | | | | | | | | ■ |
| | Leu 140 O | Tyr 41 OH | | | ■ | | | | | |
| | Arg 139 O | Ser 48 OG | ■ | ■ | | | | | | |
| | Glu 114 OE2 | Arg 49 NH1 | | | ■ | | | | | |
| | Gly 141 O | Tyr 41 OH | | | ■ | | | | | |
| | Glu 114 OE1 | Asn 105 ND2 | | | | | | ■ | | |
| Salt bridges | Glu 114 OE2 | Arg 49 NH1 | | | ■ | | | | | |
| | Glu 114 OE2 | Arg 49 NH2 | | | ■ | | | | | |

**Supplementary Table S14. Buried surface areas at the interhexameric interfaces.** PDBePISA analysis [4] was used to calculate the buried surface areas at the interhexameric interfaces (either between UL31 trimers or UL31 dimers) within the NEC-hexamer$_{AB}$, NEC-hexamer$_{CD}$, and SUP$_{UL31}$ mutant lattices. The total trimeric interface buried surface area was calculated by adding the UL31/UL31 surface areas within a trimer together. For WT HSV-1, the 4ZXS PDB structure was used.

| Trimeric interface | | | | |
|---|---|---|---|---|
| **Construct/interface** | **Chains** | | **Surface Area (Å)** | **Total surface area (Å)** |
| **WT Trimer (AB)** | **UL31$_B$/UL31$_B$/UL31$_B$** | **UL31$_B$/UL31$_B$** | 252 | 756 |
| | | **UL31$_B$/UL31$_B$** | 252 | |
| | | **UL31$_B$/UL31$_B$** | 252 | |
| **WT Trimer (CD)** | **UL31$_D$/UL31$_D$/UL31$_D$** | **UL31$_D$/UL31$_D$** | 180 | 540 |
| | | **UL31$_D$/UL31$_D$** | 180 | |
| | | **UL31$_D$/UL31$_D$** | 180 | |
| **SUP Trimer 1** | **UL31$_B$/UL31$_H$/UL31$_F$** | **UL31$_B$/UL31$_H$** | 350 | 1,117 |
| | | **UL31$_F$/UL31$_H$** | 358 | |
| | | **UL31$_B$/UL31$_F$** | 409 | |
| **SUP Trimer 2** | **UL31$_D$/UL31$_J$/UL31$_L$** | **UL31$_D$/UL31$_L$** | 357 | 804 |
| | | **UL31$_L$/UL31$_J$** | 198 | |
| | | **UL31$_J$/UL31$_D$** | 249 | |

| Dimeric Interface | | |
|---|---|---|
| **Interface** | **Construct/chains** | **Total surface area (Å)** |
| **Dimer 2** (UL31 only) | **WT UL31$_B$/UL31$_B$** | 107 |
| | **SUP UL31$_B$/UL31$_D$** | 119 |
| | **SUP UL31$_F$/UL31$_L$** | 127 |
| | **SUP UL31$_H$/UL31$_J$** | 96 |

**Supplementary Table S15. Comparison of contacts made at the WT NEC and NEC-SUP$_{UL31}$ lattice interhexameric interfaces.** Atomic contacts (hydrogen bonds or salt bridges) between heterodimers within an interhexameric lattice (shaded in blue) determined using PDBePISA [4].

| Interhexameric Contacts | | | | | | | |
|---|---|---|---|---|---|---|---|
| | Trimer | | | | | | |
| | UL31 Residue | UL31 Residue | WT A/A/A | WT D/D/D | SUP B/H/F | SUP D/J/L | |
| H-bonds | Gln 146 NE2 | Asn 149 O | | | | DL | |
| | Glu 138 OE1 | Ala 197 N | | | | | |
| | Glu 138 OE2 | Ala 197 N | | | FH | | |
| | Arg 295 NH1 | Glu 153 OE2 | | | | | |
| | Asp 286 O | Arg 155 NH2 | | | | | |
| | Asp 286 OD1 | Arg 155 NH2 | | | | | |
| | Ser 136 OG | Gly 196 N | | | BH | | |
| | Ser 136 OG | Gly 195 O | | | | JL | |
| | Arg 193 NH2 | Glu 138 OE1 | | | BF | DL | |
| | Glu 267 OE2 | Arg 155 NH1 | | | BF | | |
| | Gly 130 O | Arg 131 NE | | | BF | | |
| Salt bridge | Glu 138 OE2 | Arg 155 NH2 | | | | | |
| | Arg 295 NE | Glu 153 OE2 | | | | | |
| | Arg 295 NH1 | Glu 153 OE1 | | | | | |
| | Arg 295 NH1 | Glu 153 OE2 | | | | | |
| | Arg 295 NH2 | Glu 153 OE1 | | | | | |
| | Arg 295 NH2 | Glu 153 OE2 | | | | | |
| | Asp 286 OD1 | Arg 155 NH2 | | | | | |
| | Asp 286 OD2 | Arg 155 NH2 | | | | | |
| | Arg 193 NH2 | Glu 138 OE1 | | | BF | | |
| | Arg 193 NH2 | Glu 138 OE2 | | | | DL | |
| | Glu 267 OE2 | Arg 155 NH1 | | | BF | | |
| | Dimer 1 | | | | | | |
| | UL31 Residue | UL34 Residue | WT A/B (none) | WT C/D | SUP AB/CD (none) | SUP EF/KL | SUP GH/IJ (none) |
| H-bond | Ser 73 N | Glu 17 | | | | EL | |
| | Asp 126 O | Arg 132 NH2 | | | | | |
| | Dimer 2 | | | | | | |
| | UL31 Residue | UL31 Residue | WT B/B (none) | WT D/D (none) | SUP B/D | SUP L/F | SUP J/H (none) |
| H-bond | Asp 286 OD2 | Arg 295 NH2 | | | | | |
| | Asp 286 OD1 | Arg 295 NH2 | | | | | |
| Salt bridge | Asp 286 OD1 | Arg 295 NH2 | | | | | |
| | Asp 286 OD2 | Arg 295 NH2 | | | | | |

**Supplementary Table S16. Buried interface surface areas for the globular, hook, and the total heterodimeric interface between UL31 and UL34.** PDBePISA analysis [4] was used to calculate the buried surface areas between the globular, hook, and total heterodimer interfaces within the NEC-$WT_{AB}$, NEC-$WT_{CD}$, and the six mutant $SUP_{UL31}$ heterodimers. The globular interface area was determined by deleting the UL31 hook region (residues 54-88) prior to PDBePISA analysis. The hook interface was determined by subtracting the globular area from the total heterodimeric interface area. The total heterodimeric interface area was calculated from the entire crystal structure. For WT HSV-1, the 4ZXS PDB structure was used.

| NEC Heterodimer | Heterodimeric interface area (Å) | Globular core (Å) | Hook (Å) |
|---|---|---|---|
| **$WT_{AB}$** | 1764 | 440 | 1324 |
| **$WT_{CD}$** | 1745 | 430 | 1315 |
| **$SUP_{AB}$** | 1773 | 529 | 1244 |
| **$SUP_{CD}$** | 1780 | 467 | 1313 |
| **$SUP_{EF}$** | 1782 | 492 | 1290 |
| **$SUP_{GH}$** | 1767 | 481 | 1286 |
| **$SUP_{IJ}$** | 1742 | 498 | 1244 |
| **$SUP_{KL}$** | 1830 | 514 | 1316 |

**Supplementary Table S17. List of primers used for cloning procedures described in Materials and Methods.** All primers are listed in the 5'-3' direction. Restriction sites are underlined and mutations are bolded.

| Primer Name | Primer Sequence (5'-3') | Restriction Site |
|---|---|---|
| **Oligomeric Interface Mutants** | | |
| F252Y$_{31}$/R229L$_{31}$ and E153R$_{31}$/R229L$_{31}$ fwd A (KH199) | agcaggatcctatgacaccgacccccat | SOE/BamHI |
| F252Y$_{31}$/R229L$_{31}$ rev A (JB178) | ccggaccacgagcac**ata**cgtgctctgccacac | SOE |
| E153R$_{31}$/R229L$_{31}$ rev A (JB184) | caggaaggcgcgatgcct**gaa**tatcgtgttgatc | SOE |
| F252Y$_{31}$/R229L$_{31}$ and E153R$_{31}$/R229L$_{31}$ rev B (KH179) | aaatgcggccgcttacggcggaggaaactc | SOE/NotI |
| F252Y$_{31}$/R229L$_{31}$ fwd B (JB177) | gtgtggcagagcacg**tat**gtgctcgtggtccgg | SOE |
| E153R$_{31}$/R229L$_{31}$ fwd B (JB183) | gatcaacacgatattc**agg**catcgcgccttcctg | SOE |
| **Hexameric Interface Mutants** | | |
| K137A$_{34}$, R139A$_{34}$, and K137A$_{34}$/R139A$_{34}$ fwd A (ED058) | aaaaaagtcgacctatggcgggactgggcaagccc | SOE/SalI |
| K137A$_{34}$ rev A (ED055) | caggccgagccgcccc**cgc**gatggtgtccaggtc | SOE |
| R139A$_{34}$ rev A (ED057) | ggcatccaggccgag**cgc**cccttgatggtgtc | SOE |
| K137A$_{34}$/R139A$_{34}$ rev A (ED061) | ccgggcatccaggccgag**cgc**ccc**cgc**gatggtgtccaggtcgcc | SOE |
| K137A$_{34}$ fwd B (ED054) | gacctggacaccatc**gcg**gggcggctcggcctg | SOE |
| R139A$_{34}$ fwd B (ED056) | gacaccatcaagggg**gcg**ctcggcctggatgcc | SOE |
| K137A$_{34}$/R139A$_{34}$ fwd B (ED60) | ggcgacctggacaccatc**gcg**ggg**gcg**ctcggcctggatgcccgg | SOE |
| K137A$_{34}$, R139A$_{34}$, and K137A$_{34}$/R139A$_{34}$ rev B (ED059) | aaaaaagcggccgccttcagtcccccct | SOE/NotI |
| **Membrane Interface Mutants** | | |
| SE$_{6}$/SUP fwd (ED094) | **tg**ctcatagaccggatgctcaccg | Inverse PCR |
| SE$_{6}$/SUP fwd (ED054) | **g**gtagtgcaggtgggcggacg | Inverse PCR |
| **Crystallization Constructs** | | |
| R229L$_{31}$Δ50-306 fwd (KH178) | agcaggatcccaggagctgtgtttacac | BamHI |
| R229L$_{31}$Δ50-306 rev (KH178) | aaatgcggccgcttacggcggaggaaactc | NotI |

| | | |
|---|---|---|
| D35A$_{34}$/E37A$_{34}$ 15-185 fwd (JB36) | aaaaaagtcgacctgccttcgagggtctcgttca | SalI |
| D35A$_{34}$/E37A$_{34}$ 15-185 rev (JB162) | aaaaaagcggccgcctggcggcgcggcaca | NotI |

**Supplementary Table S18.** List of plasmids used to create the NEC constructs used in this study not previously described.

| Construct | UL31 Plasmid | UL34 Plasmid |
|---|---|---|
| *Oligomeric Interface Mutants* | | |
| NEC-$SUP_{UL31}$/$F252Y_{UL31}$ | pED20 | pJB02 |
| NEC-$SUP_{UL31}$/$E153R_{UL31}$ | pED21 | pJB02 |
| NEC-$SUP_{UL31}$/$T123Q_{UL34}$ | pJB14 | pJB89 |
| *Heterodimeric Interface Mutants* | | |
| NEC-$K137A_{UL34}$ | pKH90 | pED25 |
| NEC-$R139A_{UL34}$ | pKH90 | pED26 |
| NEC-$K137A_{UL34}$/$R139A_{UL34}$ | pKH90 | pED27 |
| NEC-$K137A_{UL34}$/$SUP_{UL31}$ | pJB14 | pED25 |
| NEC-$K137A_{UL34}$/$R139A_{UL34}$/$SUP_{UL31}$ | pJB14 | pED27 |
| *Membrane Interface Mutants* | | |
| NEC-$SE6_{UL31}$-His | pJB60 | pJB57 |
| NEC-$SUP_{UL31}$-His | pJB14 | pJB57 |
| NEC-$SUP_{UL31}$/$SE6_{UL31}$-His | pED45 | pJB57 |
